# Alterations in neural stem cell quiescence and activation in the 3xTG-AD model of Alzheimer’s Disease

**DOI:** 10.1101/2022.06.08.495344

**Authors:** Yubing Liu, Bensun C. Fong, Richard A. Harris, Marie-Michelle McNicoll, Amaal A. Abdi, Jacob B. Cuthbert, David P. Cook, Daniel Figeys, Jing Wang, Barbara C. Vanderhyden, Ruth S. Slack

## Abstract

Alzheimer’s Disease (AD) is the most common form of dementia with progressive cognitive deficits and mood disorders (Knopman et al., 2021). Recent studies have associated AD pathology with the impairment of adult neurogenesis, as indicated by impaired neural stem cell (NSCs) homeostasis (Bond et al., 2015). Recent work has further associated AD progression with a decline in the number and maturation of adult-born neurons in the SGZ, distinct from typical age-related decline (Moreno-Jiménez et al., 2019). In 3xTG-AD mice, a well-established mouse model of AD, our and other groups have demonstrated impairments to NSC pool and neural progenitor proliferation, as well as adult-born neurons, before the onset of Aβ plaques and NFTs (Hamilton et al., 2010, 2015; Rodríguez et al., 2008, 2009). However, the regulatory mechanisms underlying the functional impairment of adult NSCs remain to be resolved. Here, we employ single-cell RNA-Seq to establish population-specific defects in the 3xTG-AD mouse model during adult SGZ neurogenesis. Relative to control mice, we observe a dramatic AD-induced decrease in the primed and activated NSC population, which results in a progressive loss of cells committed to neurogenesis. Transcriptome measurements suggest that 3xTG-AD NSCs and their progeny represent enhanced ribosomal and mitochondrial biogenesis, and disturbed Notch signaling pathway. RNA velocity analysis reveals reduced NSC activation as evidenced by a large fraction of Ascl1-postive cells, instead of entering cell cycle, returning to the primed and quiescent state. This is further supported by reduced numbers of Lpar1-expressing cells, a marker of neural progenitor cells, in the SGZ. Our work explores, at a stage-specific resolution, changes in the regulatory networks guiding adult neurogenesis, and identifies niche disturbances in the regulation of NSC quiescence and activation. These NSC deficits underlying impaired neurogenesis identified in AD mice, may be key contributors underlying the compromised hippocampal function in AD.

## Introduction

Alzheimer’s Disease (AD), the most common form of dementia, is a progressive neurodegenerative disorder associated with cognitive deficits and mood disorders (Knopman et al., 2021). It is characterized most commonly by distinct neuropathological features including accumulation of β-amyloid (Aβ) plaques followed by deposition of microtubule associated protein tau (tau, also MAPT) as neurofibrillary tangles (NFTs), resulting in synaptic dysfunction, neuroinflammation and neuronal loss (Wander & Song, 2021).

Recent studies have associated AD pathology with the impairment of adult neurogenesis – the process by which a quiescent pool of neural stem cells (NSCs) give rise to adult-born neurons throughout life (Bond et al., 2015). This intricately-regulated process follows a stage-wise progression from exit from NSC quiescence, through activation to neural progenitor cell (NPC) proliferation, differentiation into neuroblasts, post-mitotic terminal maturation and subsequent integration into surrounding circuitry (Bonaguidi et al., 2016). NSCs are localized to well-established neurogenic niches in the mammalian brain, including the ventricular-subventricular zone (V-SVZ) of the lateral ventricles, as well as the subgranular zone (SGZ) of the hippocampal dentate gyrus (DG) (Gonçalves et al., 2016). SGZ NSCs generate granule cell neurons, which modify existing neural circuits by inducing neural plasticity: the process of structural reorganization that plays a role in hippocampal cognitive function, including learning, memory encoding and consolidation, and spatial navigation (Sahay et al., 2011; Toda & Gage, 2017).

In humans, the hippocampus represents one of the brain regions most damaged by AD (Coupé et al., 2019). Analysis of granule cell neurons in the DG of AD patients reveals alterations in both dendritic tree morphology and post-synaptic densities, which is similar to those observed following overexpression of tau-inducer GSK-3β (Llorens-Martín et al., 2013). Recent work has further associated AD progression with a decline in the number and maturation of adult-born DG neurons, distinct from typical age-related decline (Moreno-Jiménez et al., 2019). Together, these studies suggest an impact of AD on the generation, survival and function of adult-born neurons.

Studies in animal models have also demonstrated that AD pathology impacts adult NSC fate (Mu & Gage, 2011), including studies employing the triple-transgenic 3xTG-AD mouse model, generated by co-injecting hAPP (hAPPSwe) and hTau (hTauP301L) transgenes, under the control of the Thy1.2 promoter, into PS1M146V knock-in mouse embryos (Oddo et al., 2003). 3xTG-AD mice distinctly accumulate Aβ plaques and NFTs with age, and multiple groups have demonstrated further impairments to NSC and progenitor proliferation, as well as neurogenesis as early as 2 months of age (Hamilton et al., 2010; Rodríguez et al., 2008). Notably, these effects predate the onset of Aβ plaques and NFTs, yet are sustained as late as 18 months of age(Hamilton et al., 2010, 2015; Rodríguez et al., 2008, 2009). We have recently reported that deficits in neurogenesis can be detected as early as postnatal day 7, as indicated by reduced NSC numbers and decreased proliferation in the SGZ (McNicoll et al., in revision). Interestingly, this decrease was associated with AD-induced deregulation of oleic acid synthesis (Hamilton et al., 2015), however, the regulatory mechanisms underlying the functional impairment of adult NSCs remain to be uncovered.

Transcriptomics analyses have been useful to distinguish differential gene regulation in FACS-isolated populations obtained from the adult SVZ and SGZ (Codega et al., 2014; Than-Trong et al., 2018). Recently, single-cell transcriptomics approaches (single-cell RNASeq) have been able to provide a broad overview of the transcriptome throughout the stages of adult neurogenesis (Kalinina & Lagace, 2022), and to define the mechanisms underlying the transition between, and subtype acquisition of, distinct NSC populations (Borrett et al., 2020).

To identify the deficits underlying the impairment of neurogenesis in the 3xTG-AD model of Alzheimer’s disease, we used single-cell RNA-Seq to identify population-specific changes in the 3xTG-AD AD model during adult SGZ neurogenesis. Relative to non-transgenic (NTG) control mice, we observe a dramatic AD-induced decrease in the primed NSC population, which results in a progressive loss of cells committed to neurogenesis. Transcriptome measurements suggest that 3xTG-AD NSCs and their progeny exhibit enhanced ribosomal and mitochondrial biogenesis, along with perturbations in the Notch signaling pathway. RNA velocity analysis revealed impaired NSC activation evident by reduced cell cycle entry, and a large fraction of Ascl1-postive cells from 3xTG-AD exhibit a trend to return to a primed quiescent state. Together, our results suggest a previously-unknown phenotype for AD pathology, affecting adult neurogenesis at early stages involving defects in NSC activation, that precede the subsequent impairments in neuronal differentiation. Thus, we reveal, at a stage-specific resolution, changes in the regulatory networks guiding adult neurogenesis, and identify niche disturbances in the regulation of NSC quiescence and activation as a key deficit of AD.

## Results

### Reduced adult neurogenesis in the 3xTG-AD Brain

The 3xTG-AD mouse model of Alzheimer’s Disease exhibits significant deficits in adult neurogenesis together with a progressive cognitive decline (Belfiore et al., 2019; Hamilton et al., 2010). Prior reports have observed impairments to the NSC pool in AD patients, consistent with those observed in the mouse model of 3xTG-AD (Hamilton et al., 2015; Moreno-Jiménez et al., 2019; Tobin et al., 2019). Here, we used a tamoxifen (Tam)-inducible Nestin-CreER^T2^;ROSA26-EYFP Cre-reporter mouse line that was cross-bred with 3xTG-AD or NTG control mice (Cicero et al., 2009; Zhu et al., 2012), targeting the adult-born NSC and their progeny. Prior to single-cell transcriptomic analysis we performed immunostaining with cell type specific markers to confirm the impairment of neurogenesis previously described in this animal model. SRY (Sex-determining region Y)-box 2 (Sox2), a marker of NSCs and regulator of NSC homeostasis(Julian et al., 2015; Sarkar & Hochedlinger, 2013), was significantly reduced in 3xTG-AD at 4-weeks post TAM injection (**Fig.1A**). We also show a reduction in the HOP homeobox (Hopx) gene expression in the quiescent NSC population of the SGZ (Berg et al., 2019; Li et al., 2015; Shin et al., 2015) (**Fig. 1B**). To validate the previously reported reduction in progenitor proliferation in the SGZ, we measured EdU incorporation (**Fig. 1B**). The 3xTG-AD mice exhibited decreased proliferation, as evidenced by a reduction in the number of EdU positive cells and EdU+/YFP+ double-labeled cells in the SGZ. Consistent with reduced proliferation there was also a decrease in the number of newly generated neuroblasts, measured by Doublecortin (Dcx) (Brown et al., 2003). Thus, the number of proliferating cells (EdU), the new-born neurons (DCX), as well as the Dcx/YFP double positive population, were significantly reduced in the 3xTG-AD SGZ (**Fig. 1B**). Taken together, our initial characterization confirms the presence of deficits in the NSC population and adult neurogenesis in 3xTG-AD mice at 3 months of age, consistent with previously reported findings (Hamilton et al., 2010; Rodríguez et al., 2008). To elucidate the mechanisms underlying the deficit in neurogenesis in this AD model, we used a single-cell transcriptomic approach to characterize the transcriptional program within distinct cell populations.

**Figure 1:**
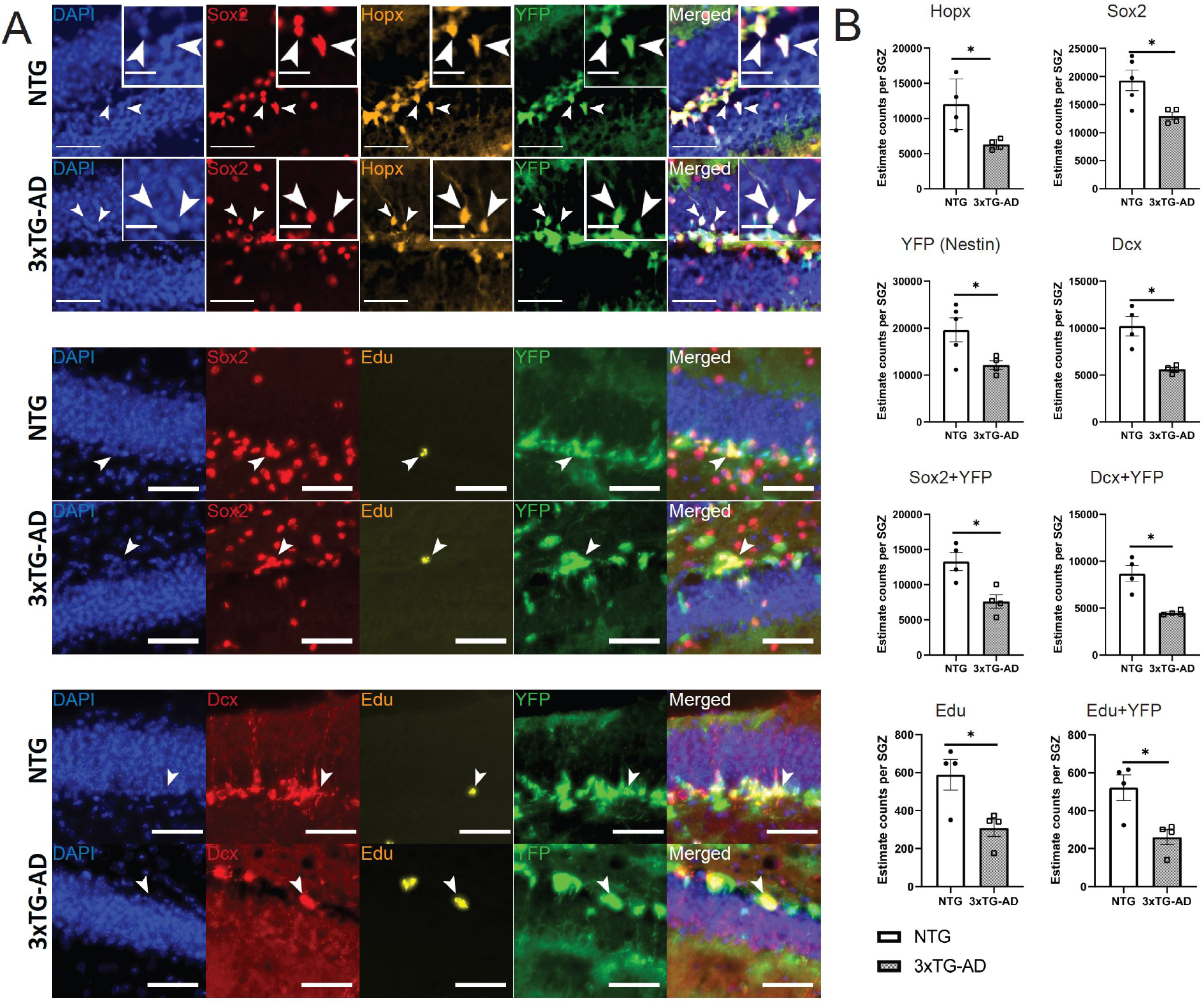
Impaired neurogenesis in the SGZ of 3xTG-AD mice. (A)Immunostaining of neural stem cell markers (Sox2, Hopx, and recombined YFP), and newborn neuron marker Dcx, and proliferation marker EdU in the SGZ of NTG and 3xTG-AD brains, at three months of age. Scale bar, 25μm. Insets represent magnified views. Arrowheads point to cells with colocalization. Scale bar, 10 μm. (B)Quantification of neural stem cell markers (Sox2, Hopx, and recombined YFP), and newborn neuron marker Dcx, and proliferation marker EdU in the SGZ of NTG and 3xTG-AD brains, at three months of age. Unpaired and two-tailed Student t-test. * p < 0.05. n =4.

### Single Cell Transcriptome of the Nestin-expressing populations in the SGZ

Using the Tam inducible Nestin-CreER^T2^ bred to the ROSA26 YFP reporter mice, we isolated YFP expressing cells at 4 weeks post Tam directly from the adult SGZ by fluorescence-activated cell sorting (FACS), and then performed single cell RNA-seq (**Fig. 2A**). The transcriptomes of 11154 NTG and 8646 3xTG-AD cells were independently analyzed using the 10X Genomics platform. Following quality control and detection of YFP expression (**Fig. S1**), 6105 NTG and 5160 3xTG-AD cells were recovered for further analysis. The NTG and 3xTG-AD transcriptomes were integrated using Seurat, and UMAP for unsupervised clustering based on gene expression. Ten major tissue-resident cell types were identified (**Fig. 2B**). The neurogenic lineage consists of five cell populations including cluster 0 – NSCs enriched for astrocyte markers (astrocyte-rich NSC), cluster 1 - quiescent NSCs (qNSC), cluster 2 - NSCs primed for activation (pNSC), cluster 3 – transiently-amplifying progenitors (TAPs), and cluster 4 – neuroblast and new-born neurons (neurons). In addition, there were cluster 5 - oligodendrocyte, cluster 6 - Mural cells, cluster 7 - Endothelial cells, cluster 8 - Ependymal cells, and cluster 9 - oligodendrocyte progenitor cells (OPC) (**Fig. 2B**). A heatmap demonstrating both distinct and continuous expression of the top ten overrepresented genes in each cluster (**Fig. 2C**) was used for clustering validation. Individual gene expression heatmaps of NTG and 3xTG-AD cells were plotted to compare their cell cluster-specific markers and revealed similar pattern (**Fig. S2**). For non-neurogenic clusters, Mog, Mobp and Mag were used as markers of oligodendrocytes; Gpr17, Tnr and C1ql1 for oligodendrocyte progenitor cells (Marques et al., 2019); Myl9, Hspb1 and Myo1b shared by mural cells and endothelial cells; Flt1, Ly6c1 and Cldn5 for endothelial cells (He et al., 2016; Zhao et al., 2018); and enriched expression of Ccdc153, Rsph1 and Meig1 were found in ependymal cells (Mizrak et al., 2019).

**Figure 2:**
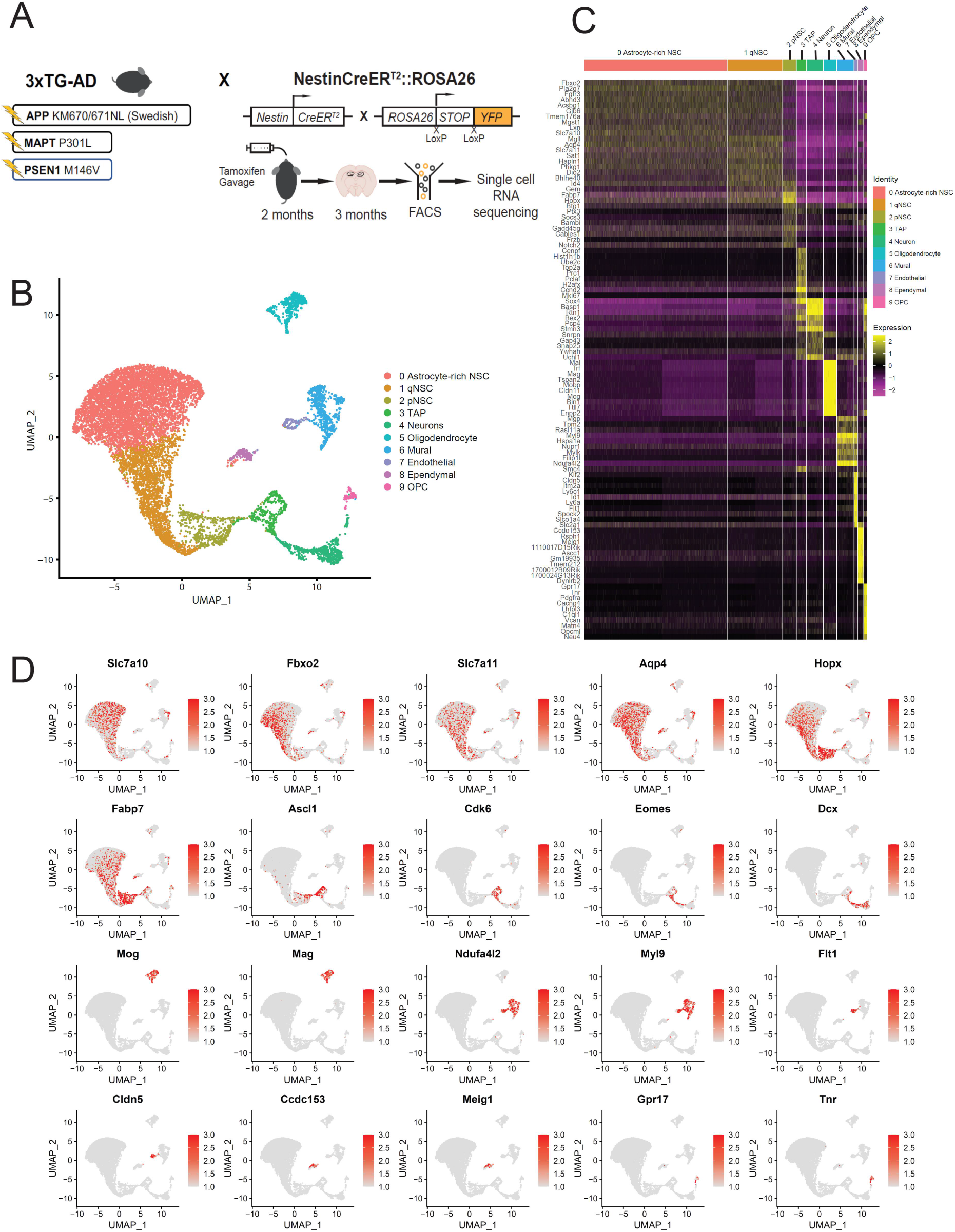
Ten cell clusters with recombined YFP expression identified in the SGZ. (A)Experimental design. Inducible NestinCreER^T2^;R26-EYFP mice were crossed with either NTG or 3xTG-AD mice. Tamoxifen induction occurred at 2 months of age and hippocampal cells were isolated at 3 months of age. YFP-positive cells were isolated using fluorescence-activated cell sorting (FACS) and subjected to single cell RNA-seq. (B)Integrated UMAP projection of NTG and 3xTG-AD cells, separated using Seurat into ten cell clusters based on their expression profiles. (C)Heatmap demonstrating genes representative of each cluster, reflecting cluster-specific expression. (D)UMAP projections demonstrating curated genes representative of each cluster, reflecting cluster-specific expression.

The YFP-expressing cells generated by Tam treatment of the Nestin-CreER^T2^;ROSA26-EYFP animals, were found in five cell clusters expressing characteristic stem, progenitor, neurogenic and neuronal markers, consistent with previous reports (**Fig. 2C-D**) (Bernal & Arranz, 2018; Youssef et al., 2018). The majority of cells having stem cell characteristics were represented in clusters 0, 1 and 2: 0-astrocyte-rich NSCs, 1-qNSCs and 2 -pNSCs (**Fig. 2C-D, S3 and S4**). The astrocyte-rich NSC cluster 0 shows enriched expression of astrocyte markers, including Slc7a10 and Fbxo2 (Batiuk et al., 2020) (**Fig. 2D, S3C and S4A**). Quiescence markers Aqp4, and Id4 were highly expressed in the cluster 1 qNSC, and downregulated in the cluster 2 pNSC and differentiating populations (clusters 3 and 4), consistent with prior characterization (Artegiani et al., 2017; Shin et al., 2015) (**Fig. 2D and S3B**). Dbi and Sfrp1, found in primed NSCs, were expressed in our cluster 2 pNSC (Basak et al., 2018) (**Fig. 2D and S3A**). The expression of the NPC marker, Fabp7 (Fatty acid binding protein 7) was dramatically induced in the cluster 2 pNSC (Li et al., 2015; Matsumata et al., 2012) (**Fig. 2C-D**). Using gene ontology, relative to the cluster 0 astrocyte-rich, the cluster 1 qNSC was enriched in genes for “nervous system development” and “cell population proliferation” (**Fig. S5**). Relative to the cluster 1 qNSC, genes involved in “translation” and “cell population proliferation” were enriched as cells transition to the ‘primed’ state, found in the cluster 2 pNSC (**Fig. S5**). These findings support the heterogeneity of NSCs in the hippocampus and identify a set of markers for the progressive NSC transition between stages of quiescence, activation and differentiation.

The expression of Ascl1 and Eomes (Tbr2) further identify the gradual transition from activated NSCs in the cluster 2 pNSC to the cluster 3 TAPs; a fraction of TAPs are in M-phase, demonstrating expression of cell cycle-related genes including Cdk6, Mki67 and Top2a (**Fig. 2C-D**). Finally, the expression of differentiation markers Dcx and Stmn3 are induced in neuroblasts from the cluster 4 Neurons, consistent with the acquisition of the immature neuronal phenotype (**Fig. 2C-D**). The Nestin-CreER^T2^-labeled population four weeks post-Tam, exhibited integrated clusters advancing from quiescence towards activation, proliferation, and differentiation, and revealed a stagespecific transcriptome signature over the course of adult neurogenesis, consistent with previous reports (Shin et al., 2015).

### Changes in NSC Homeostasis in the 3xTG-AD SGZ

To better characterize cluster-specific differences in the 3xTG-AD mice, we examined our samples within the integrated UMAP projection for all the cell clusters (**Fig. S6A**), focusing on the five neurogenic cell clusters (0-4) (**Fig. 3A**). Although each transcriptome reflected a similar-sized neurogenic population (5302 cells in NTG, 4314 in 3xTG-AD), there were significantly decreased proportions of pNSCs (Cluster 2, - 41.5%), TAP (Cluster 3, −17.9%) and neuroblasts/neurons (Cluster 4, −28.3%), and a correspondingly increased proportion of the qNSC (Cluster 1, 14%) in the 3xTG-AD relative to the NTG control (**Figs. 3B-C, S6B-C**). Cluster-specific differential gene expression was established between NTG and 3xTG-AD samples. While many important developmental genes were similar between NTG and 3xTG-AD samples, genes associated with translation and mitochondrial function were significantly enriched in the 3xTG-AD cells (**Figs. 3D-F, S8**). Of the significantly differentially-expressed genes, those upregulated in response to 3xTG-AD were induced in the clusters 2 and 3 (pNSC and TAP), including mt-Co2, mt-Nd1 and multiple ribosomal genes (**Fig. 3D-F**), reflecting a general increase in energy demand associated with activation and proliferation in cell derived from the animal models. This suggests the possible compensatory upregulation of translation machinery and mitochondrial biogenesis, both of which are tightly controlled to maintain cell homeostasis (Gabut et al., 2020; Khacho et al., 2019; Tahmasebi et al., 2019).

**Figure 3:**
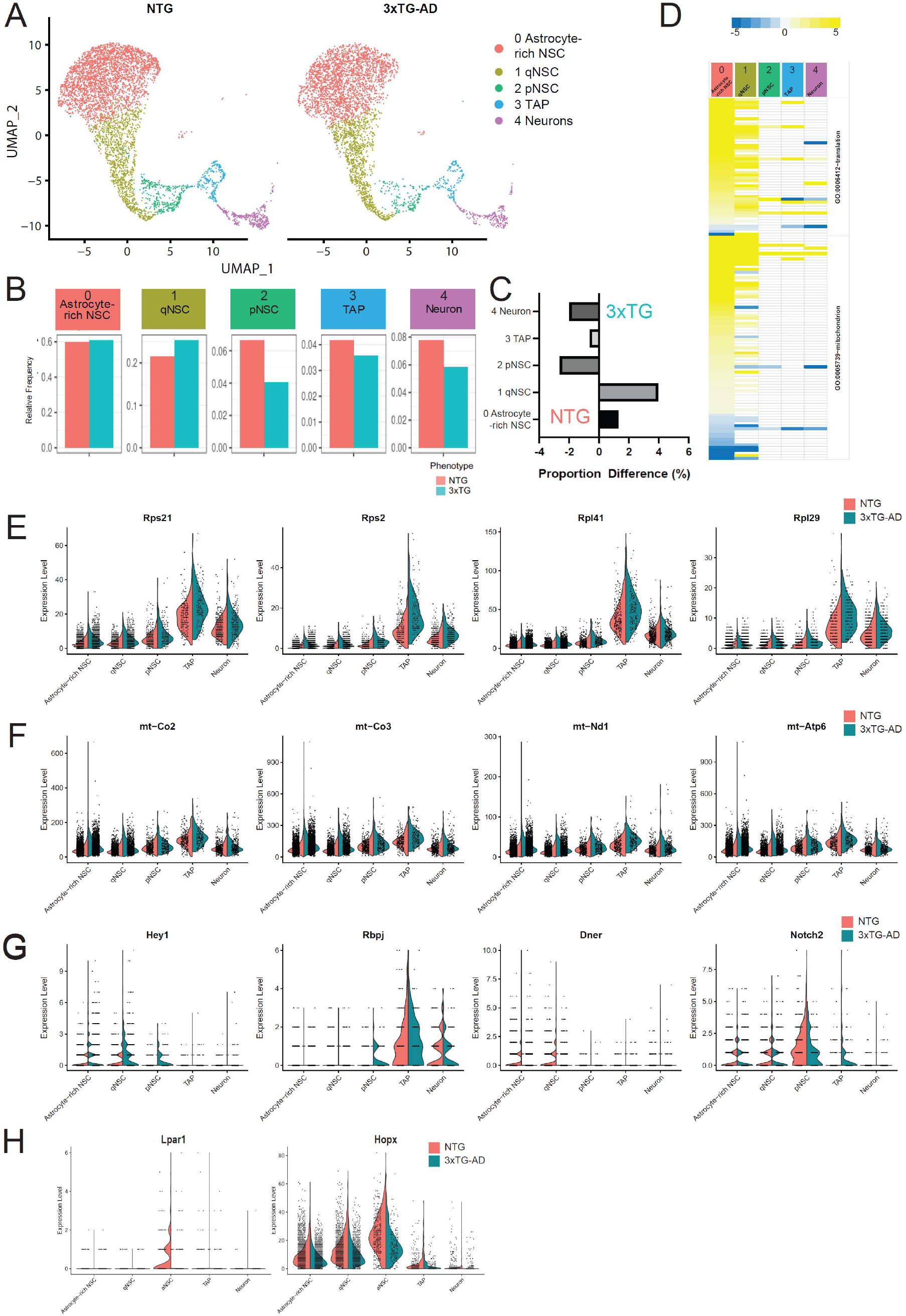
Single-cell RNA-seq reveals intrinsic defects in 3xTG-AD NSCs. (A)UMAP projections separating NTG and 3xTG-AD samples for the neurogenic lineage. (B)Cluster proportion for each sample, expressed as a percentage of total cell number within the neurogenic clusters. (C)Proportional difference between NTG and 3xTG-AD cluster proportions from B. Positive indicates a greater proportion in the 3xTG-AD sample, negative indicates a greater proportion in the NTG sample. (D)Gene ontology analysis reflecting the expression fold change of indicated gene sets. Yellow reflects upregulation, blue reflects downregulation in 3xTG-AD compared to NTG. Violin plots demonstrating expression of (E) representative ribosomal genes, (F) mitochondrial genes, (G) genes implicated in the Notch signaling pathway, and (H) Lpar1 and Hopx, separated by cluster and sample.

Consistent with impaired NSC activation, several genes in the Notch signaling pathway, that play a critical role in NSC homeostasis, were dysregulated in the 3xTG-AD cells (**Fig. 3G**). For example, Hey1 is a Notch downstream effector that plays a critical role in promoting return to quiescence of slowly dividing NPC (Harada et al., 2021). Rbpj is a key intracellular signal mediator of the Notch pathway and is required for NSC maintenance in undifferentiated state (Imayoshi et al., 2010). Both Hey1 and Rbpj exhibited aberrantly elevated expression in the cluster 2 pNSC of the 3xTG-AD SGZ (**Fig. 3G**), consistent with maintenance of quiescence (increased proportion of the cluster 1 qNSC in the 3xTG-AD SGZ). These findings suggest an imbalance in the regulation of NSC quiescence and activation in cells isolated from the SGZ neurogenic niche of 3xTG-AD animals.

Our data also revealed an impairment in Lpar1 expression in the cluster 2 pNSC in the 3xTG AD model, which also suggested an impairment in NSC activation (**Fig. 3H**). Lpar1 (Lysophosphatidic acid receptor 1) is a receptor of lysophosphatidic acid, a natural bioactive lipid mediator, and has been shown as a marker of neural precursors (Walker et al., 2016). In the absence of Lpar1, cell proliferation in the dentate gyrus is reduced (Matas-Rico et al., 2008). In the wild type, NTG transcriptome, Lpar1 expression is low in the astrocyte-enriched and qNSC clusters, and becomes upregulated in the pNSC cluster, consistent with a role for Lpar1 in NSC activation (Walker et al., 2016) (**Fig. 3H**). However, this upregulation of Lpar1 was absent in the 3xTG-AD cells (**Fig. 3H**), which in line with the observation of reduced proliferation in the 3xTG-AD SGZ (**Fig. 1**). Similarly, Hopx and Fabp7, of which expression increases as cells undergo activation (Li et al., 2015; Matsumata et al., 2012), also exhibited reduced expression in the cluster 2 pNSC of 3xTG-AD cells (**Fig. 3H and S9**). Taken together, these data suggest an impairment in the regulation of adult NSC quiescence and activation in the 3xTG-AD SGZ niche.

### RNA Velocity Reveals Defective Activation in 3xTG-AD NSCs

To better characterize the potential defect in NSC activation suggested by the differential gene expression in single cell RNA-seq, we used RNA velocity to predict the transcriptional fate of cell populations based on RNA processing. RNA velocity measures relative differences in splicing kinetics (Bergen et al., 2020; La Manno et al., 2018), enabling us to predict the fate transitions of 3xTG-AD and NTG cells during the course of adult neurogenesis. We focused our analysis on the clusters 2, 3 and 4 (pNSC, TAP and neuroblast/neuron), to establish a linear trend regarding NSC fate (**Fig. 4A**). The NTG velocity stream suggests the continuous progression of the cluster 2 pNSCs through activation, proliferation and differentiation. Consistent with the dynamic transition between the clusters 2 and 3 (pNSC and TAPs), a few cells in the TAP cluster appears destined to return towards the pNSC fate, possibly the self-renewing population, with the vast majority predicted to progress toward differentiation into neurons (**Fig. 4B**). However, though the differentiation program appeared consistent between the clusters 3 and 4 (TAP and neuroblast/neuron) in both NTG and 3xTG-AD, we observed a dramatic shift in the fate of the cluster 2 pNSC cells in the 3xTG-AD transcriptome, suggesting that pNSCs fail to transition towards the expected TAP fate but instead are destined to return to a more primed stem cell state (**Fig. 4B-C**). As RNA velocity studies also suggested an impairment in NSC activation, we asked whether characteristic genes regulating this transition may be differentially regulated in the cluster 2 pNSC of the 3xTG-AD transcriptome. RNA velocity in individual genes reflects cluster-specific trends in their instantaneous regulation trend (Bergen et al., 2020). We observed significant changes in the RNA processing dynamics in the Notch signaling pathway, including Hey1 and Rbpj, and as well as Lpar1 (**Fig. 3G and 4D**). The upregulation of Hey1 and Rbpj in the cluster 2 pNSC and their aberrant mRNA dynamics support the interpretation of impaired activation in the 3xTG-AD SGZ. RNA velocity also reveals aberrant mRNA dynamics of Lpar1 in the clusters 2, 3 and 4 (pNSC, TAP and neuroblast/neuron). Our results showed that the induction of Lpar1 in the cluster 2 pNSC of NTG cells (shown in green) becomes repressed in 3xTG-AD pNSCs (shown in red), consistent with reduced expression of Lpar1 in the 3xTG-AD SGZ (**Fig. 4D**). Thus, RNA dynamics further supports the increased return to quiescence revealed by the aberrant regulation of the Notch pathway (Hey1, Rbpj), and absence of Lpar1 induction in the cluster 2 pNSCs, supporting the hypothesis that the activation of adult NSCs is impaired leading to an overall reduction in adult neurogenesis in the 3xTG-AD mice.

**Figure 4:**
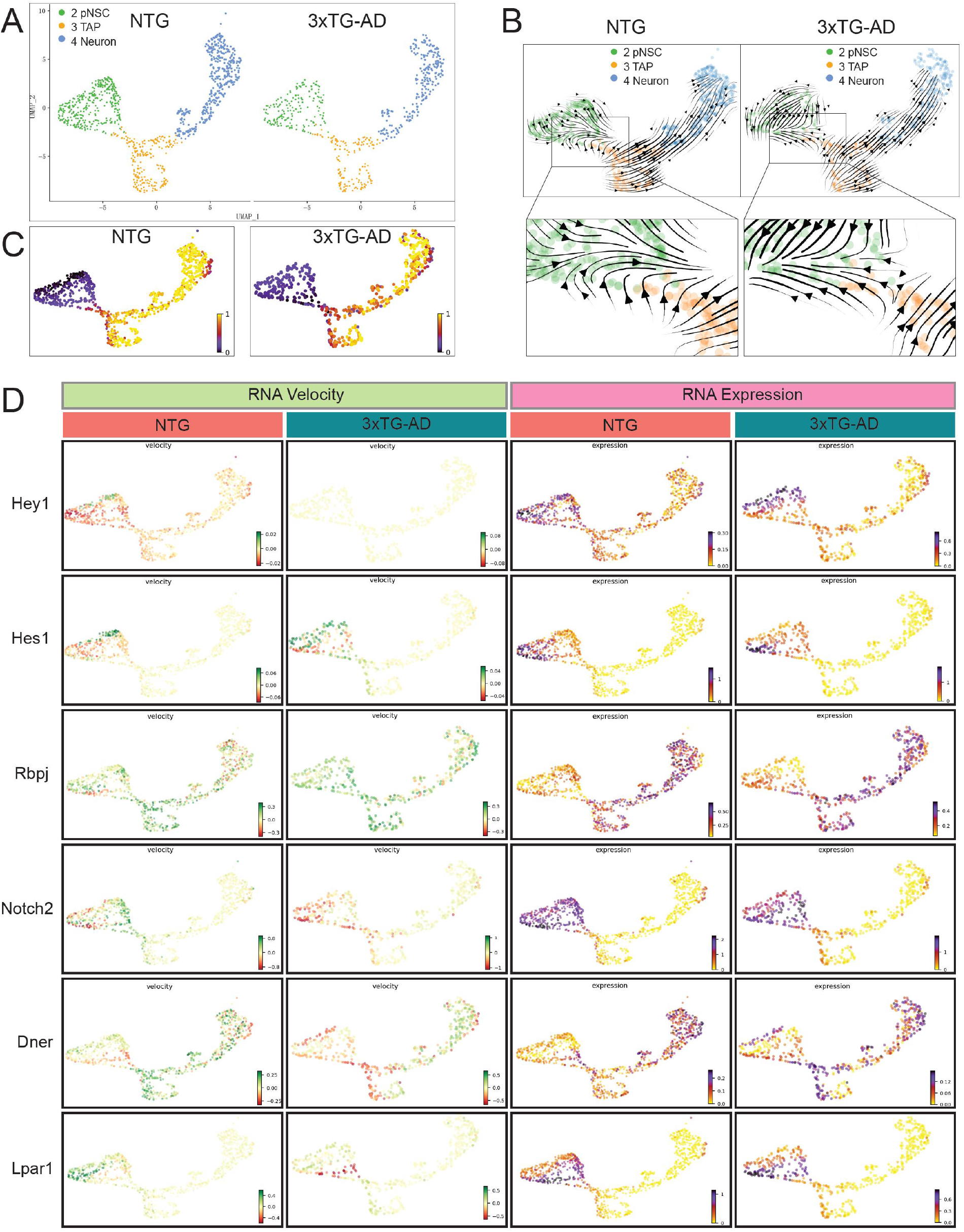
RNA velocity identifies defects in NSC activation. (A)UMAP projections isolating pNSC, TAP and neuron clusters, by sample. (B)RNA velocity analysis of projections from A, by sample. Lower panels are magnified views of transition between pNSC and TAP cells. (C)RNA pseudo-time analysis of projections from A. (D)RNA velocity and RNA expression plots for genes implicated in the Notch signaling pathway (Hey1, Hes1, Rbpj, Notch2 and Dner) and Lpar1.

To validate the changes identified by single cell RNA-seq and RNA velocity, *in vivo* immunostaining of Lpar1 was performed on both NTG and 3xTG-AD brains (**Fig. 5A**). Compared to the NTG control, a significantly reduced number of Lpar1 and Sox2/Lpar1 expressing cells were present in the 3xTG-AD SGZ (**Fig. 5B-D**). Thus, the reduced Lpar1 expression at mRNA level and elevated expression of Hey1 and Rbpj in the cluster 2 pNSC strongly support a defect in NSC activation in the neurogenic niche of the 3xTG animal model of AD.

**Figure 5:**
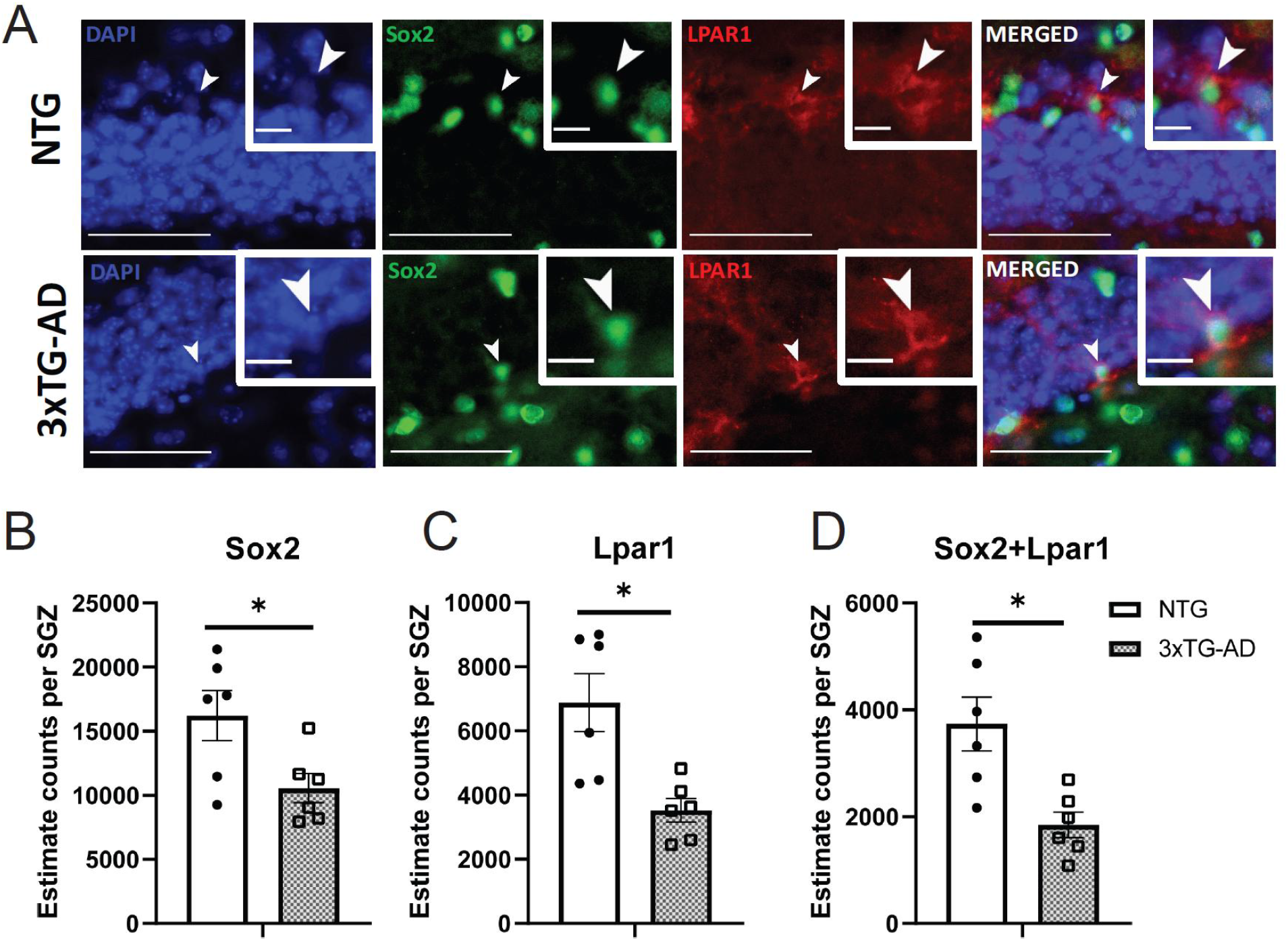
NSC activation markers are compromised in 3xTG-AD. (A) Immunostaining of Sox2 and Lpar1 in the SGZ of NTG and 3xTG-AD brains. Quantification of Sox2-expression (B), Lpar1-expressing (C), and Sox2/Lpar1 co-expressing (D) cells in the SGZ of NTG and 3xTG-AD brains. Unpaired and twotailed Student t-test. * *p* < 0.05. n = 6.

## Discussion

This study provides a number of important insights into the dysregulation of NSC function in the SGZ niche of AD mice. First, we provide a single cell analysis of the Tam-inducible Nestin-YFP expressing cells sorted directly from the SGZ niche in the adult 3XTG AD model along with age- and genetic background-matched control animals. Second, we report significant differences in the distribution of cells among clusters from the quiescent NSCs over the course of adult neurogenesis. We find that 3xTG-AD mice possess a greater proportion of qNSC cells, along with a corresponding reduction of pNSCs and activated transit amplifying cells. Third, we identify changes in gene expression profiles consistent with impairments in NSC activation, including elevated expression of Hey1 and Rbpj, and the absence of Lpar1 induction in the cluster 2 pNSC that are primed for activation. Third, RNA velocity measurements also indicate an impairment in NSC activation, and reveal increased numbers of cells reverting back to the quiescence state. Together our studies reveal an impairment in the regulation of NSC quiescence and activation in the SGZ of the 3xTG-AD mouse model.

Single cell RNA-seq has provided molecular signatures of different cell types at different developmental stages in many tissues, including NSCs in the brain (Kalinina & Lagace, 2022; Shin et al., 2015). During adult neurogenesis, activated and proliferating NSCs have distinct molecular signatures compared to qNSCs, such as the induction of Ascl1 and Tbr2 gene expression (**Fig. 2D**). However, qNSCs and astrocytes share a set of common molecular markers, including Slc1a3, Gfap and Aldoc. Previous studies examined the transcriptome of isolated astrocytes and qNSC through FACS based on a combination of cell surface markers (GFAP-GFP^+^PROM1^−^EGFR^−^ for astrocytes and GFAP-GFP^+^PROM1^+^EGFR^−^ for qNSCs), revealed that astrocyte and qNSCs exhibited similar expression patterns (Dulken et al., 2017). Here, we study all Nestin-YFP-positive NSCs and their progeny. The cluster 0 astrocyte-enriched NSC and cluster 1 qNSC share a combination of astrocyte and NSC markers, consistent with previous studies (Batiuk et al., 2020)(Shin et al., 2015). Borrett et al. identified a set of astrocyte and dormant NSC markers, which are enriched in our clusters 0 and 1 (Borrett et al., 2020) (**Fig. S3**). Similarly, Basak et al. provided a gene set of qNSC and pNSC markers (Basak et al., 2018), which exhibited enriched expression in our cluster 0, 1 and 2. Primed NSCs (pNSC) represents the population prior to activation, with elevated expression of ribosomal and mitochondrial genes (Fig. 3D-F), which we find is proportionately reduced in the SGZ of AD mice. Our study reveals the changes in the molecular signatures of adult NSCs and their progeny in the 3xTG AD mouse model.

Our single cell RNA-seq reveal significant changes over the course of adult neurogenesis beginning with quiescent NSCs in 3xTG-AD mice. These findings are consistent with previous studies which revealed compromised neurogenesis in the 3xTG-AD SGZ niche (Hamilton et al., 2015; Rodríguez et al., 2008). Here we confirm impaired adult neurogenesis in the 3xTG-AD SGZ, and we report significant changes associated with defective NSC activation. Relative to NTG controls, we observed that 3xTG-AD mice exhibit a proportionally greater qNSC population (cluster 1, 14% increase), a reduction in the proportion of primed (pre-activation stage) NSCs in cluster 2 (Lpar1-expressing pNSC cells, −41.5%), cluster 3 (Ascl1-expressing TAP, −17.9%) and cluster 4 (Dcx-expressing newborn neurons, −28.3%) (**Fig. 3A-C**). These cell counts revealing a proportional increase in the quiescent population together with a reduction in activated NSCs, suggest a defect in NSC activation in AD mice.

This analysis highlights a number of genes and pathways, the dysregulation of which, may be involved in the impaired activation of NSC in 3xTG-AD SGZ niche. The Notch signaling pathway plays a critical role in NSC homeostasis in the brain and perturbations in Notch signaling could have a direct impact on the regulation of NSC activation and quiescence (Bjornson et al., 2012; Imayoshi et al., 2010). We report elevated expression of Hey1 and Rbpj in the pNSC population of 3xTG-AD, of which a proportion of these cells failed to progress into proliferation (**Figs. 3G, 4C**). In fact, RNA velocity analysis also revealed aberrant expression dynamics of Hey1 and Rbpj in this population together with an increased reversion to the quiescent state. In addition, Dner, the Delta/notch-like epidermal growth factor (EGF)-related receptor (Hsieh et al., 2013) exhibited reduced expression 3xTG-AD pathology in the NSC mice. Notch2 which was shown to regulate NSC quiescence and proliferation (Engler et al., 2018; R. Zhang et al., 2019), was also mis-regulated in the pNSC and TAP population of 3xTG-AD cells (**Fig. 3G and 4D**). The mechanisms underlying the dysregulation of the Notch signaling pathway in the NSCs of the 3xTG-AD SGZ, and the impact on impaired NSC activation, will be the subject of future studies.

Here, we applied scVelo on NTG and 3xTG-AD cells to study the RNA dynamics of individual genes, including RNA transcription, splicing and degradation, in cells transitioning from quiescence to differentiation (Bergen et al., 2020). scVelo is a powerful tool to identify difference in RNA regulation in cells in transient states during developmental processes. The RNA dynamics in these cell populations strongly support our observation of impaired NSC activation. We identified a proportion of Ascl1-expressing TAP cells (cluster 3) from 3xTG-AD SGZ returning to a primed state (cluster 2, pNSC), which is consistent with the findings revealing a increase in the cluster 1 population (qNSC) and a corresponding reduction clusters 2 to 4 (pNSC, TAP and neuron) in the 3xTG-AD SGZ. The RNA dynamics of individual genes further supports these findings. Aberrant induction of Hey1 and Rbpj is consistent with their elevated expression in the cluster 2 pNSC cells in 3xTG-AD; and the aberrant repression of Lpar1, an activation marker, reflects its reduced expression in the same cluster, consistent with impaired NSC activation.

Lipid metabolism has been shown to regulate the homeostasis of NSC. The Jessberger group reported that metabolic shifts of fatty acid oxidation regulate NSC activity, and de novo lipogenesis had a profound effect on NSPC quiescence and proliferation in the hippocampus (Knobloch et al., 2013, 2017). Hamilton et al. reported aberrant accumulation of oleic acid-containing lipid drops in the 3xTG-AD SVZ, which inhibits NSC proliferation (Hamilton et al., 2015). As a receptor of LPA (lysophosphatidic acid), Lpar1 is expressed in the neural progenitors of SGZ and it is required for their proliferation (Matas-Rico et al., 2008). Here we reported decreased Lpar1 expression in the pNSC population (cluster 2), which was also corroborated by RNA dynamics which also revealed impaired induction of Lpar1 in the cluster 2 pNSC cells. This observation was further validated by *in vivo* immunostaining, suggesting that aberrant Lpar1 regulation may contribute to the impaired NSC activation in our AD mouse model.

In conclusion, we report the previously-unknown finding of impaired regulation of adult NSC quiescence and activation of adult NSCs occurring in the hippocampus of 3xTG-AD mice. Exploring the mechanisms underlying this imbalance in adult NSC homeostasis may provide new therapeutic perspectives in AD.

## Methods

### Animals

The triple-transgenic mouse mice (3xTG-AD) were purchased from the Jackson Laboratory and previously generated at the University of California Irvine, USA (Oddo et al., 2003). Under the control of the same mouse Thy1.2 regulatory element, two transgenes carrying human familial mutations (human APP^/Swe^ and human tau^/P301L^) were integrated into the same locus of homozygous PSEN1^/M146V^ knock-in mice (JAX stock number: 004807). Wild type mice from the same genetic background were served as non-transgenic control (NTG, JAX stock number: 101045).

The *Nestin-CreER^T2^* mouse line (Nestin-Cre) was a gift from Dr. Suzanne J Baker at the University of Tennessee Health Science Center (Cicero et al., 2009; Zhu et al., 2012). *Nestin-CreER^T2^* mice bear the *CreER^T2^* transgene that is driven by the *Nestin* promoter and an enhancer in the second intron, an internal ribosomal entry size and human placental alkaline phosphatase (*hPLAP)* (Zimmerman et al., 1994). Cre-activity was mapped through cross-breeding with the *ROSA26YFP* mice to generate *Nestin-CreER^T2^;ROSA26YFP* mice (Srinivas et al., 2001). For single cell RNA-seq experiments, 3xTG-AD or NTG mice were cross-breed with *Nestin-CreER^T2^;ROSA26YFP* mice to generate mice with conditional YFP labeling of Nestin-positive cells for lineage tracing *3xTG;Nestin-CreER^T2^;ROSA26YFP or NTG;Nestin-CreER^T2^;ROSA26YFP*. All experimental protocols were performed in compliance with University of Ottawa policy concerning the care and use of animals in research (protocol #ME-1765). Genotyping primer sequences are shown in Table S3.

### Tamoxifen Administration

*Cre recombination was induced by administration of tamoxifen (TAM) through oral gavage at a dose of 200mg/kg in NTG;Nestin-CreER^T2^;ROSA26YFP or 3xTG;Nestin-CreER^T2^;ROSA26YFP* mice daily at 2 months of age for five consecutive days (Khacho et al., 2016).

### Animal Euthanasia

Animals were euthanized with intraperitoneal injection of Euthanyl (Sodium Pentobarbital) at 120 mg/kg. To collect brains for live cells, brains were then dissected and the SGZ or SVZ was collected for experiments. To fix the brains for immunostaining experiments, a perfusion needle was inserted into the left ventricle and the right atrium was open for systemic drainage. Mice were flushed with 20 ml of 0.9% cold saline and then perfused with 20 ml of 4% cold paraformaldehyde (PFA) (pH 7.4) to fix. Brains were dissected and post-fixed in 4% PFA at 4°C for 24 hours. Brains were then incubated and stored in 20% sucrose in PBS at 4°C with supplement of 0.3% sodium azide until processed further.

### Single Cell Collection and Fluorescence-activated cell sorting (FACS)

SGZ was dissected coronally in ice-cold ACSF under a Leica M165 FC stereomicroscope from *NTG;Nestin-CreER^T2^;ROSA26YFP or 3xTG;Nestin-CreER^T2^;ROSA26YFP* mice. Tissue was collected in 1.5ml Eppendorf tube with ice-cold ACSF, transferred to pre-warmed papain solution at 20 U/ml (Worthington Cat. PAP3126), homogenized with a pestle and immediately digested for 10 mins after homogenized with a pestle (Corning, PES-15-B-SI) at 37°C on a rotator. An equal volume of resuspension medium that contains 0.5 mg/ml of DNase I (Roche Cat. # 11284932001) and 10% of fetal bovine serum (Wisent Cat. # 080150) was added to the digested tissue in papain solution. Tissue was triturated 4 to 6 times with P1000 micropipet and incubated for 5 mins at room temperature. Cell suspension was collected in 15 ml tubes and cell pellets were triturated again with another 1 ml of resuspension medium. Cell suspension from different trituration was pooled and the final volume was brought to 3.9 ml. 1.1 ml of 90% Percoll in PBS was added to 3.9 ml of cell suspension. After gentle mixing, cells were span down at 500 xg for 12.5 mins at 4°C. Cell pellet was washed with DMEM:F12 and then resuspended in sorting medium (DMEM:F12 without phenol red, Invitrogen Life Tech. Cat. # 21041-025, plus 1% BSA). YFP-positive cells from 2-3 mice of each genotype were collected after viability screening with propidium iodide staining (ThermoFisher, Cat. # P3566) on a Beckman Coulter MoFlo platform in the Flow Cytometry and Cell Sorting Facility located in the Ottawa Hospital Sprott Center (Ottawa, Canada).

### Single Cell RNA TrpNSCriptome (scRNA-seq)

Freshly collected YFP-positive cells were sent to StemCore Laboratories located in the Ottawa Hospital Sprott Center, for both *NTG;Nestin-CreER^T2^;ROSA26YFP and 3xTG;Nestin-CreER^T2^;ROSA26YFP* samples. The trpNSCriptome of viable single cells was analyzed using the 10x Genomics Chromium Single Cell Assay platform, individually for each genotype. A single cell RNA library was created following the single cell 3’ Reagent Kits v2 user guide (Cat: CG0052, 10x Genomics) on the Chromium Single Cell Instrument (10x Genomics) using Single Cell 3’ Library & Gel Bead Kit v2 and Chip Kit (P/N 120236 and P/N 120237, 10x Genomics). The cDNA library was then purified with SPRIselect (Beckman Coulter). Agilent Fragment Analyzer was used to evaluate the size distribution and yield (Agilent). The sequencing was performed on Illumina NextSeq 500 instrument (Illumina) using the following parameters 28 bp Read1, 8 bp I7 Index, 0 bp I5 Index, and 98 bp Read2. Single-end reads were aligned to the reference genome, mm10, using the CellRanger pipeline (10X Genomics), after demultiplexing cell barcodes and UMI (unique molecular identifiers) barcodes. The output cell-gene matrix contains UMI counts by genes and by cell barcodes. Gene expression profiles were then mapped using Seurat v3 in R studio (Stuart et al., 2019).

### Single Cell Selection and Clustering

Cells that meet the following criteria were selected for further analysis: 1) with more than 600 genes recorded; 2) with less than 7500 genes recorded; 3) with mitochondria content less than 20% of total RNA counts; and 4) with detectable YFP expression. RNA reads were normalized with the function of sctransform “SCT”. Based on normalized RNA reads, cells were clustered using the following parameters: 1) dims =1:8; and 2) resolution = 0.2. Ten cell clusters were identified based on normalized expression profiles of each cell. Based on the original RNA reads, cluster-specific markers were calculated with the function of FindAllMarkers. Differential expression between NTG and 3xTG-AD for each cluster was analyzed with the function of FindMarkers. Heatmaps of cluster-specific genes were mapped on normalized data. FeaturePlot (Umap) and VlnPlot (violin plot) of specific genes were mapped on original RNA reads.

Cell clusters with characteristics of the neurogenic lineage were sub-grouped to show differential expression of selected genes. Activated NSC, TAP and Neurons were sub-grouped and reclustered using the following parameters: 1) dims =1:13; and 2) resolution = 0.1. RNA velocity data were generated in Seurat based on normalized data, and analyzed in Python-based Spyder program using the “deterministic” mode (La Manno et al., 2018; Stuart et al., 2019).

### Tissue Processing for Immunostaining

Each brain was cut into two hemispheres. One half hemisphere was frozen in - 40°C isopentane (Thermo Fisher Scientific) for 60 seconds and then preserved in aluminum foil on dry ice or in −80°C freezers until cryosection. Serial coronal sectioning at 30μm thickness were performed to reveal the whole SGZ using the Leica CM1850. SGZ sections were collected sequentially from the beginning to the end of the dentate gyrus into 9 wells of a 24-well plate containing PBS and 0.01% sodium azide at 4°C.

### Immunostaining

For immunofluorescence staining, sections in one of the nine wells from each brain were rinsed 3 times in wash buffer (PBS) for 5 minutes and then permeabilized 3 times with incubation buffer (0.1% Triton X-100 and 0.1% Tween-20 in PBS) for 5 minutes. Sections were incubated overnight with primary antibodies in incubation buffer at 4°C (Hopx, 1:1000, HPA030180, Atlas Antibodies; Lpar1, 1:1000, NBP1-03363, Novus Biologicals; Sox2, 1:2000, GT15098, Neuromics; YFP, 1:1000). Sections were rinsed 3 times in wash buffer for 5 minutes, and then incubated with secondary antibodies and 4’, 6-diamidino-2-phenylindole (DAPI) in incubation buffer for 2 hours at room temperature (anti-rabbit-Cy3, 1:1000, 711-165-152, Jackson; anti-mouse Alexa488, 1:1000, 715-165-151, Jackson; anti-goat Cy5, 1:1000, 705-165-147, Jackson; DAPI, 0.1 μg/ml, D9542, Sigma-Aldrich). Sections were mounted with Immunomount (Genetex) on FisherbrandTM SuperfrostTM Plus Microscope Slides (Thermo Fisher Scientific). Immunofluorescence images were taken with DeltaVision Elite-Olympus IX-71 or ZEISS Axioscan 7 at a 20X magnification. Images were then processed and quantified with Fiji (Image J).

### Cell Counting and Statistical Analysis

Cells expressing the marker of interest were quantified along the entire SGZ of the DG. The number of all positive cells was multiplied by 9 to obtain the estimated total cell number of the whole DG (Khacho et al., 2016). Two-tailed and unpaired student’s t-test was used for statistical analysis using GraphPad PRISM software (GraphPad Software, Inc).

