## Supplemental figures for "Alterations in neural stem cell quiescence and activation in the 3xTG-AD model of Alzheimer’s Disease": Liu Fong 06-08 Supplementary.pdf

### Supplementary Figure Legends

**Figure S1: Quality control of single cell RNA-seq**

A) Quality of single cell RNA sequencing and removal criteria of bad quality cells for downstream analysis
B) Distribution of RNA counts and RNA features. The expression level of mitochondria content (mt-), ribosomal proteins (Rpl- and Rps-), and YFP expression.
C) Scatter plots of RNA counts and RNA features, mitochondria content (mt-) and RNA features, ribosomal proteins (Rps-) and RNA features, YFP expression level and RNA features

A

| | NTG | 3xTG-AD |
|---|---|---|
| Estimated Number of Cells | 11,154 | 8,646 |
| Fraction Reads in Cells | 93.40% | 93.20% |
| Mean Reads per Cell | 43,418 | 53,624 |
| Median Genes per Cell | 2,344 | 2,367 |
| Total Genes Detected | 17,466 | 17,298 |
| Median UMI Counts per Cell | 5,070 | 5,404 |
| Number of Cells<br>nFeature RNA > 600<br>nFeature RNA < 7500<br>percent.yfp > 0<br>percent.mt < 20 | 6,105 | 5,160 |

B

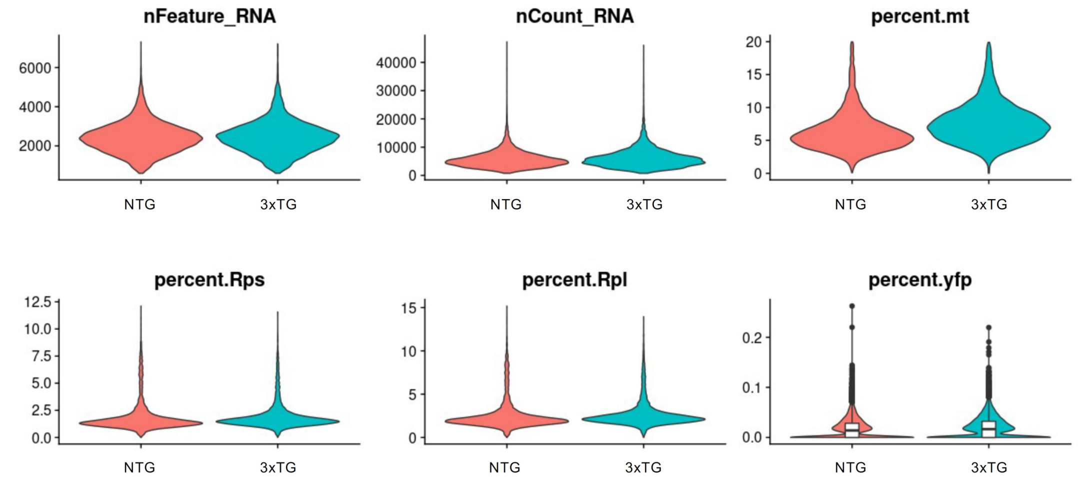


C

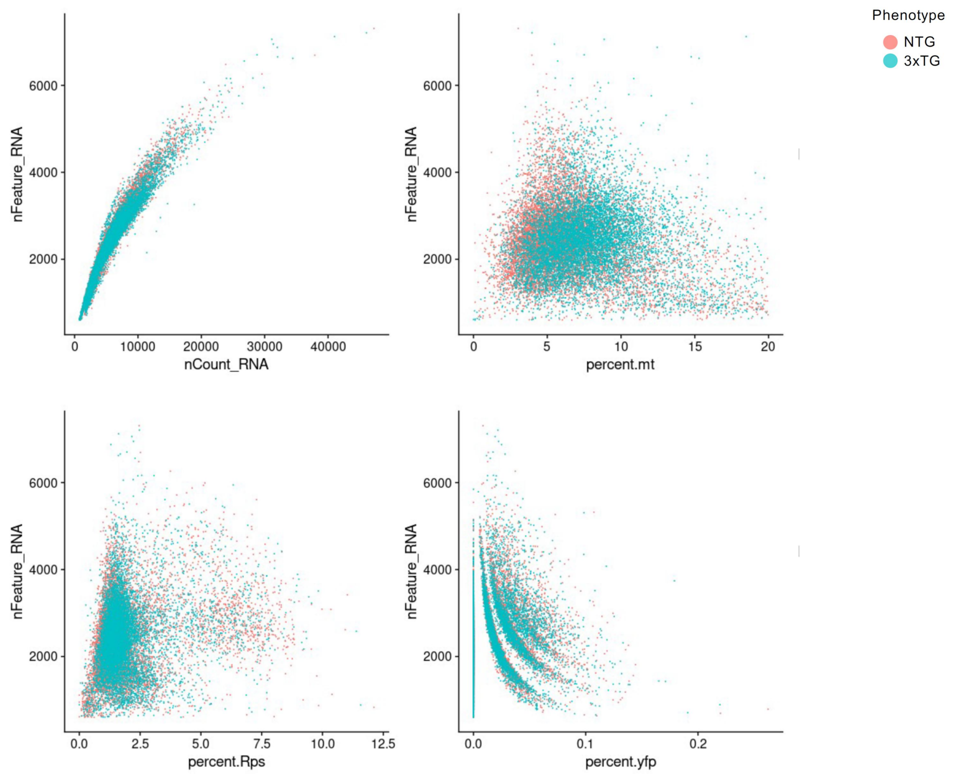

**Figure S2: Cell cluster-specific gene expression**

Ten representative genes in each cluster were selected to show cluster-specific scaled RNA expression as Heatmap for NTG (A) and for 3xTG-AD data (B).

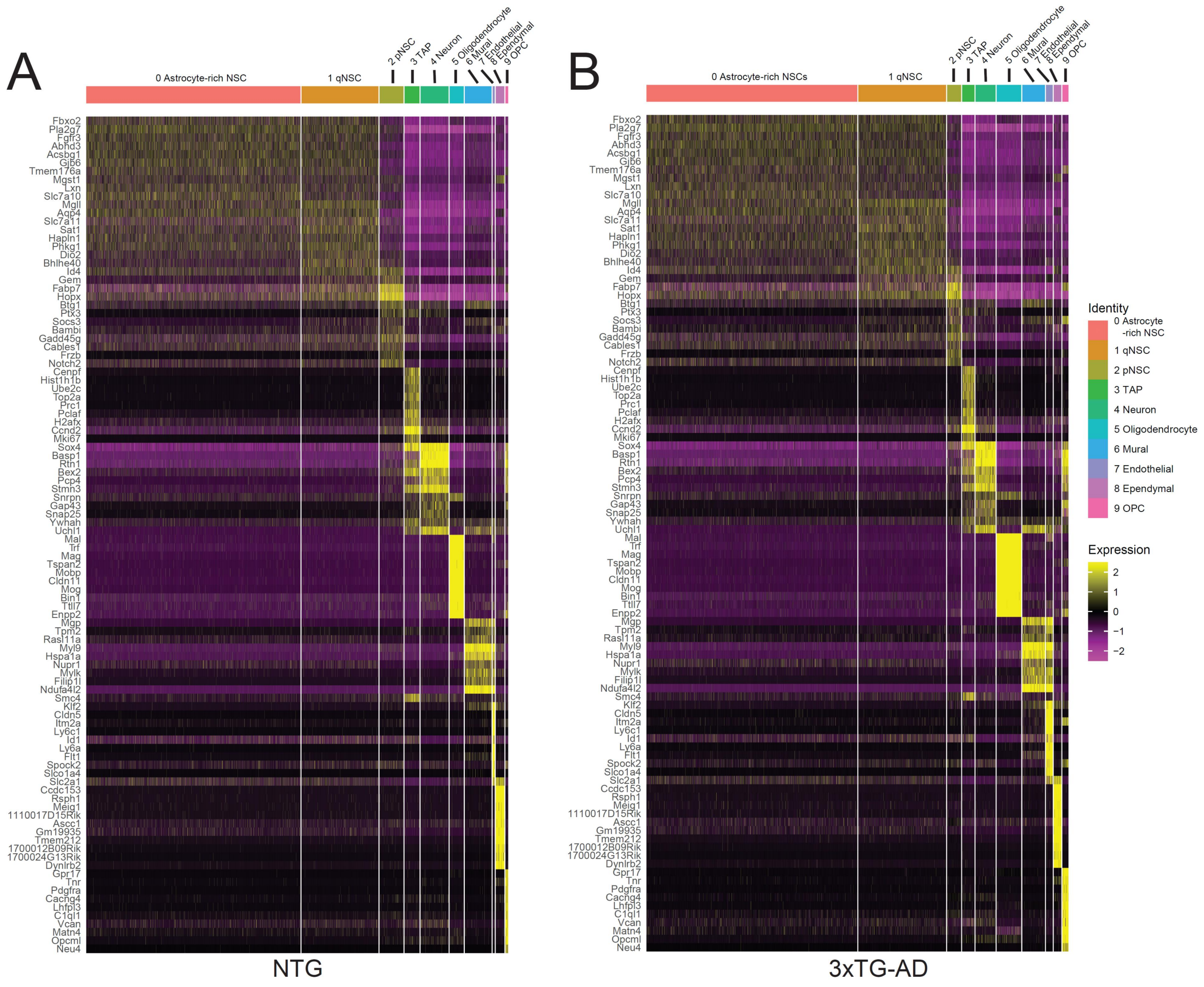
A
0 Astrocyte-rich NSC
1 qNSC
2 pNSC
3 TAP
4 Neuron
5 Oligodendrocyte
6 Mural
7 Endothelial
8 Ependymal
9 OPC
Fbxo2
Pla2g7
Fgfr3
Abhd3
Acsbg1
Gjb6
Tmem176a
Mgst1
Lxn
Slc7a10
Mgll
Aqp4
Slc7a11
Sat1
Hapln1
Phkg1
Dio2
Bhlhe40
Id4
Gem
Fabp7
Hopx
Btg1
Ptx3
Socs3
Bambi
Gadd45g
Cables1
Frzb
Notch2
Cenpf
Hist1h1b
Ube2c
Top2a
Prc1
Pclaf
H2afx
Ccnd2
Mki67
Sox4
Basp1
Rtn1
Bex2
Pcp4
Stmn3
Snrpn
Gap43
Snap25
Ywhah
Uchl1
Mal
Trf
Mag
Tspan2
Mobp
Cldn11
Mog
Bin1
Ttll7
Enpp2
Mgp
Tpm2
Rasl11a
Myl9
Hspa1a
Nupr1
Mylk
Filip1l
Ndufa4l2
Smc4
Klf2
Cldn5
Itm2a
Ly6c1
Id1
Ly6a
Flt1
Spock2
Slco1a4
Slc2a1
Ccdc153
Rsph1
Meig1
1110017D15Rik
Ascc1
Gm19935
Tmem212
1700012B09Rik
1700024G13Rik
Dynlrb2
Gpr17
Tnr
Pdgfra
Cacng4
Lhfpl3
C1ql1
Vcan
Matn4
Opcml
Neu4
NTG
B
0 Astrocyte-rich NSCs
1 qNSC
2 pNSC
3 TAP
4 Neuron
5 Oligodendrocyte
6 Mural
7 Endothelial
8 Ependymal
9 OPC
Fbxo2
Pla2g7
Fgfr3
Abhd3
Acsbg1
Gjb6
Tmem176a
Mgst1
Lxn
Slc7a10
Mgll
Aqp4
Slc7a11
Sat1
Hapln1
Phkg1
Dio2
Bhlhe40
Id4
Gem
Fabp7
Hopx
Btg1
Ptx3
Socs3
Bambi
Gadd45g
Cables1
Frzb
Notch2
Cenpf
Hist1h1b
Ube2c
Top2a
Prc1
Pclaf
H2afx
Ccnd2
Mki67
Sox4
Basp1
Rtn1
Bex2
Pcp4
Stmn3
Snrpn
Gap43
Snap25
Ywhah
Uchl1
Mal
Trf
Mag
Tspan2
Mobp
Cldn11
Mog
Bin1
Ttll7
Enpp2
Mgp
Tpm2
Rasl11a
Myl9
Hspa1a
Nupr1
Mylk
Filip1l
Ndufa4l2
Smc4
Klf2
Cldn5
Itm2a
Ly6c1
Id1
Ly6a
Flt1
Spock2
Slco1a4
Slc2a1
Ccdc153
Rsph1
Meig1
1110017D15Rik
Ascc1
Gm19935
Tmem212
1700012B09Rik
1700024G13Rik
Dynlrb2
Gpr17
Tnr
Pdgfra
Cacng4
Lhfpl3
C1ql1
Vcan
Matn4
Opcml
Neu4
3xTG-AD
Identity
0 Astrocyte -rich NSC
1 qNSC
2 pNSC
3 TAP
4 Neuron
5 Oligodendrocyte
6 Mural
7 Endothelial
8 Ependymal
9 OPC
Expression
2
1
0
−1
−2

**Figure S3: The expression of reported cell markers in our data**

(A) Cell markers for qNSC, pNSC and aNSC (Basak et al., 2018)

(B) Reported qNSC markers (Shin et al., 2015; Artegiani et al., 2017; Llorens-Bobadill et al., 2015; Urban, Blomfield and Guillemot review 2019)

(C) Astrocyte markers (Batiuk et al., 2020).

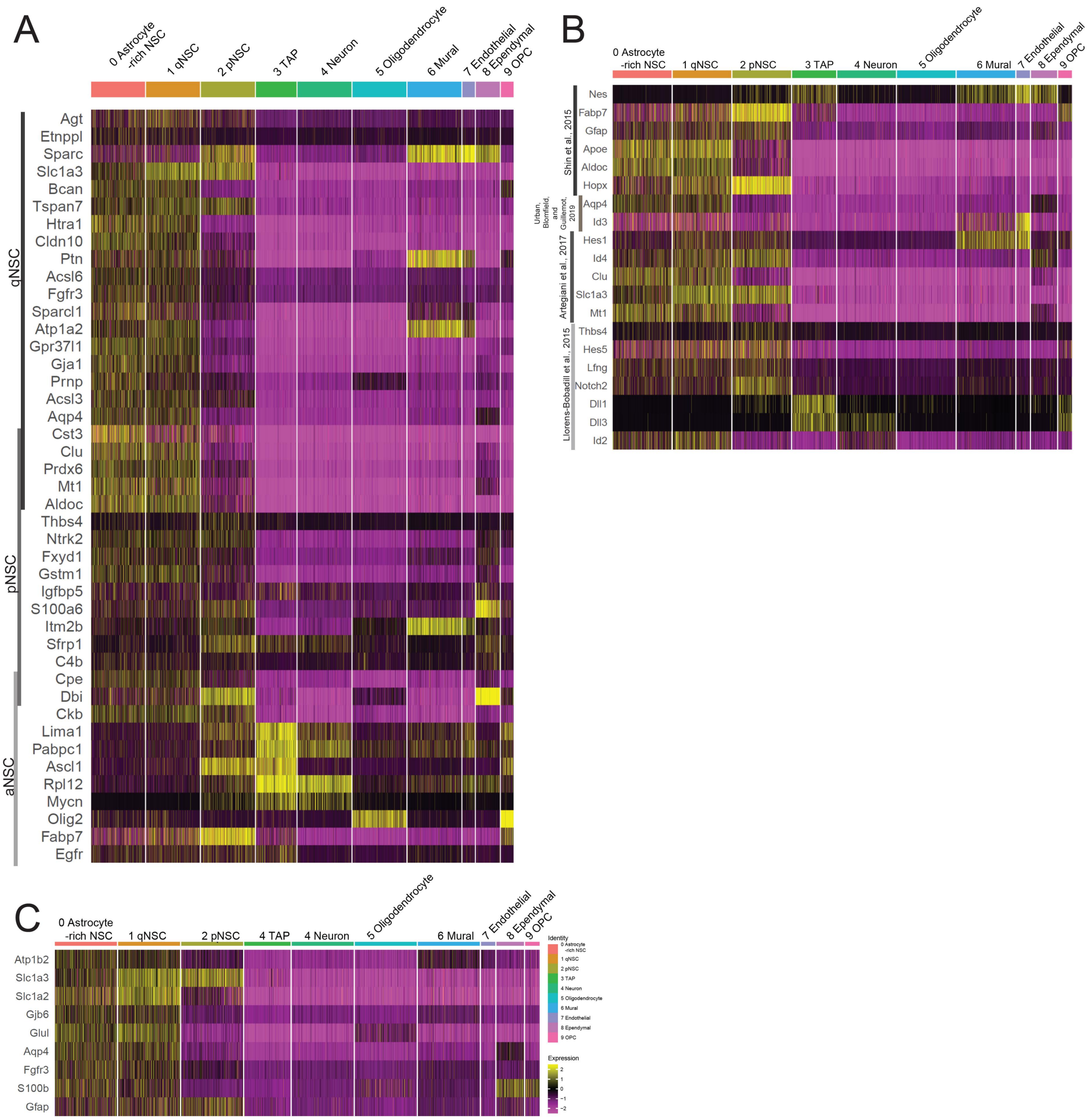

A
0 Astrocyte -rich NSC
1 qNSC
2 pNSC
3 TAP
4 Neuron
5 Oligodendrocyte
6 Mural
7 Endothelial
8 Ependymal
9 OPC
qNSC
pNSC
aNSC
Agt
Etnppl
Sparc
Slc1a3
Bcan
Tspan7
Htra1
Cldn10
Ptn
Acsl6
Fgfr3
Sparcl1
Atp1a2
Gpr37l1
Gja1
Prnp
Acsl3
Aqp4
Cst3
Clu
Prdx6
Mt1
Aldoc
Thbs4
Ntrk2
Fxyd1
Gstm1
Igfbp5
S100a6
Itm2b
Sfrp1
C4b
Cpe
Dbi
Ckb
Lima1
Pabpc1
Ascl1
Rpl12
Mycn
Olig2
Fabp7
Egfr
B
0 Astrocyte -rich NSC
1 qNSC
2 pNSC
3 TAP
4 Neuron
5 Oligodendrocyte
6 Mural
7 Endothelial
8 Ependymal
9 OPC
Shin et al., 2015
Urban, Blomfield, and Guillemot, 2019
Artegiani et al., 2017
Llorens-Bobadill et al., 2015
Nes
Fabp7
Gfap
Apoe
Aldoc
Hopx
Aqp4
Id3
Hes1
Id4
Clu
Slc1a3
Mt1
Thbs4
Hes5
Lfng
Notch2
Dll1
Dll3
Id2
C
0 Astrocyte -rich NSC
1 qNSC
2 pNSC
4 TAP
4 Neuron
5 Oligodendrocyte
6 Mural
7 Endothelial
8 Ependymal
9 OPC
Atp1b2
Slc1a3
Slc1a2
Gjb6
Glul
Aqp4
Fgfr3
S100b
Gfap
Identity
Expression
2
1
0
-1
-2


(D)

**Figure S4: Feature plots of reported cell markers**

(A) Astrocyte markers (Batiuk et al., 2020).

(B) Reported qNSC markers (Shin et al., 2015; Artegiani et al., 2017; Llorens-Bobadill et al., 2015; Urban, Blomfield and Guillemot review 2019).

A

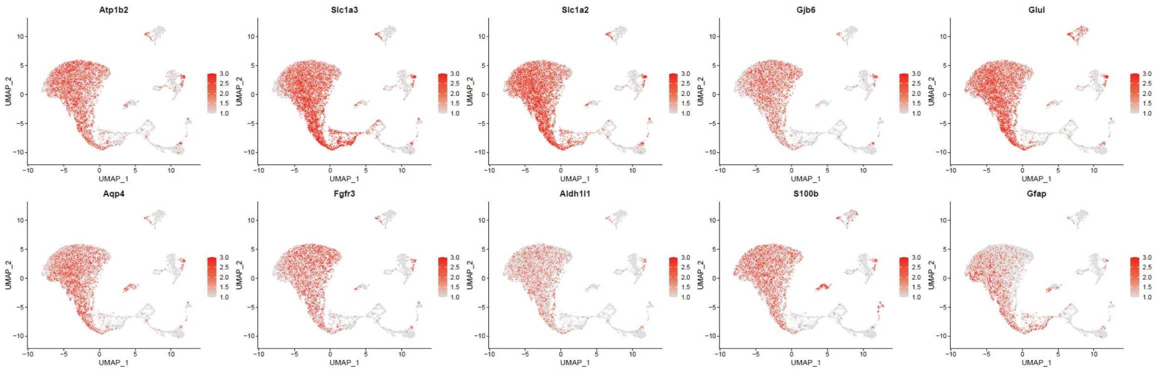
Atp1b2
Slc1a3
Slc1a2
Gjb6
Glul
Aqp4
Fgfr3
Aldh1l1
S100b
Gfap
UMAP_1
UMAP_2

B

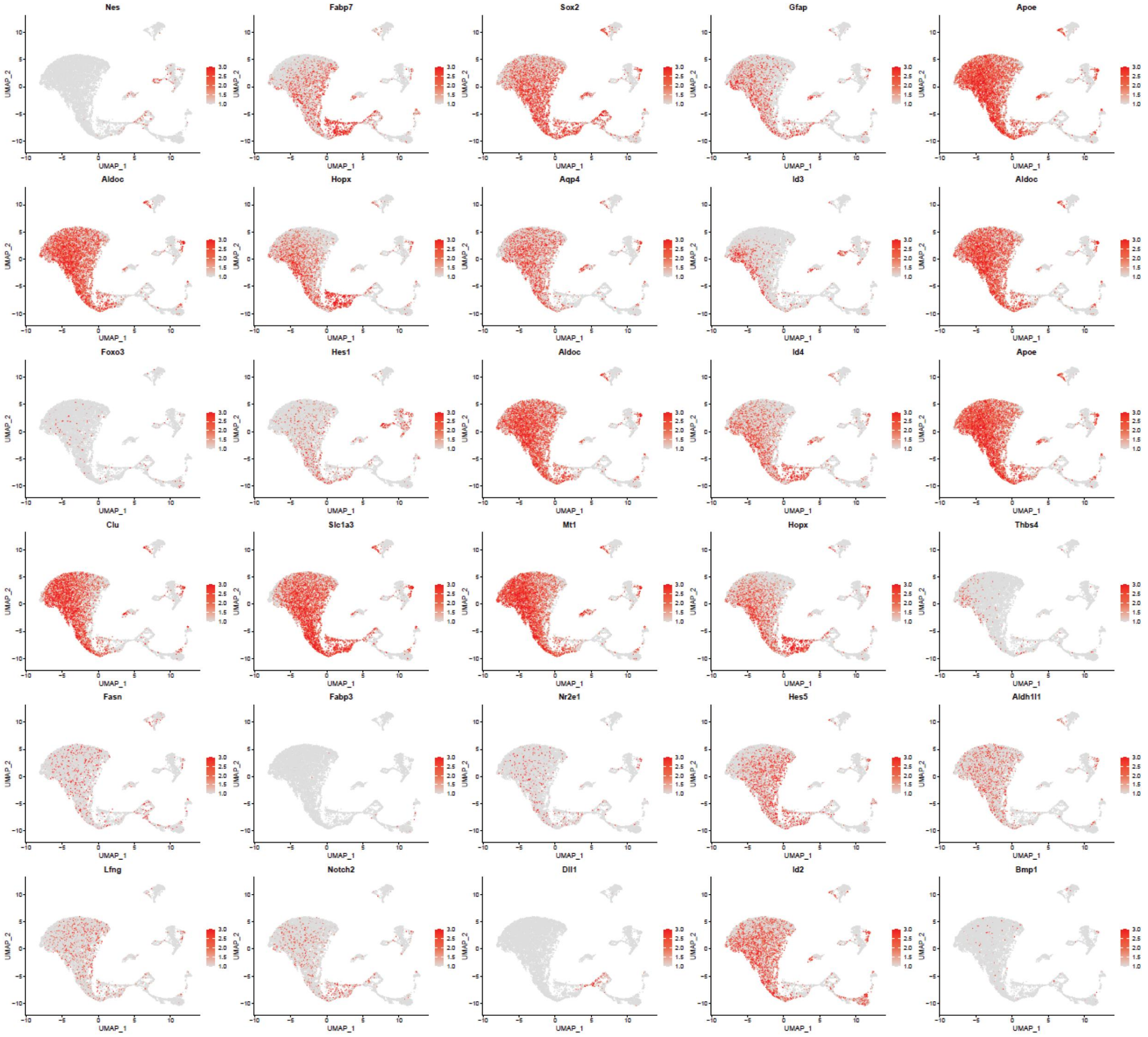
Nes
Fabp7
Sox2
Gfap
Apoe
Aldoc
Hopx
Aqp4
Id3
Aldoc
Foxo3
Hes1
Aldoc
Id4
Apoe
Clu
Slc1a3
Mt1
Hopx
Thbs4
Fasn
Fabp3
Nr2e1
Hes5
Aldh1l1
Lfng
Notch2
Dll1
Id2
Bmp1
UMAP_1
UMAP_2

**Figure S5: Gene ontology analysis of enriched genes in NSCs population**

(A) Gene ontology of qNSC-enriched genes over dNSCs
(B) Gene ontology of pNSC-enriched genes over qNSCs

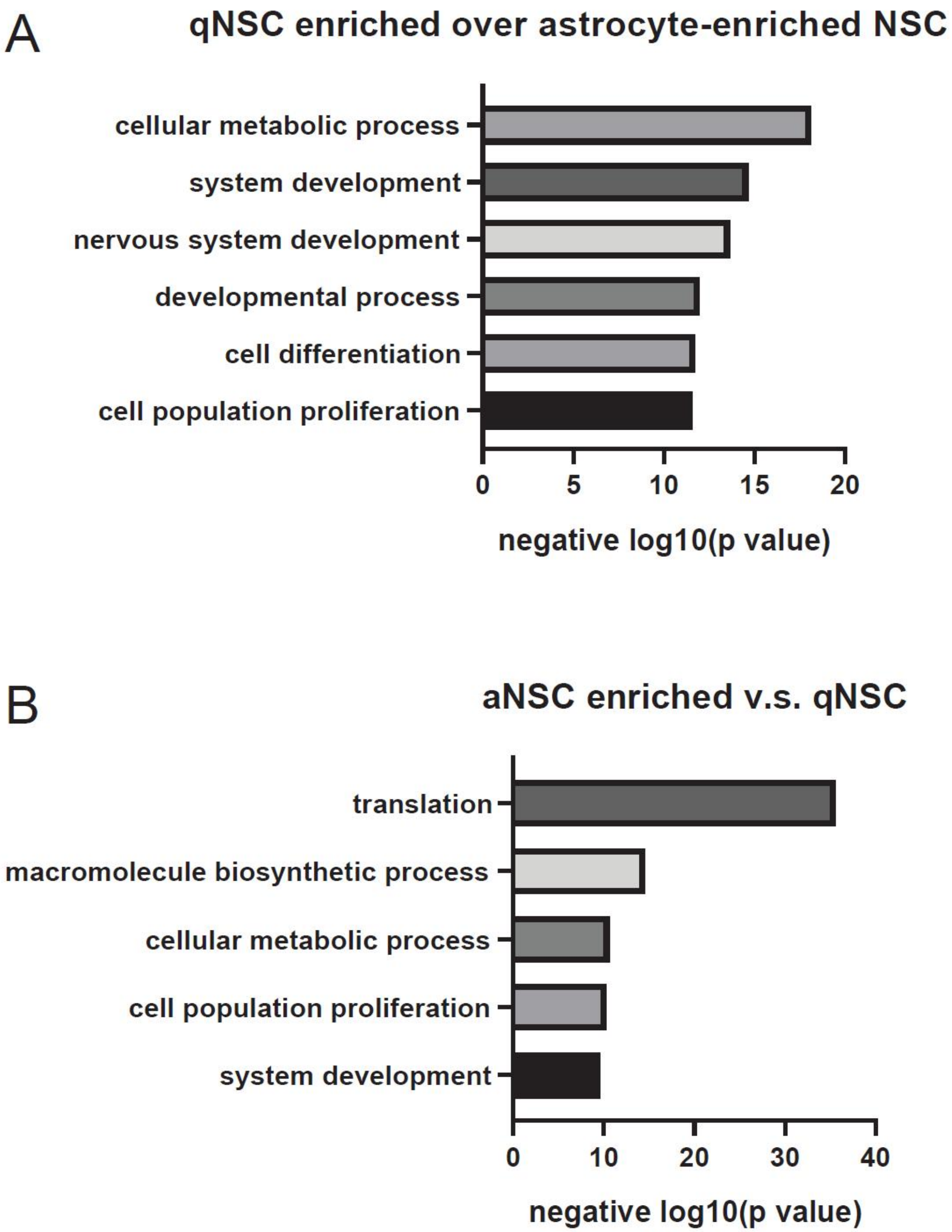
A
qNSC enriched over astrocyte-enriched NSC
cellular metabolic process
system development
nervous system development
developmental process
cell differentiation
cell population proliferation
0
5
10
15
20
negative log10(p value)
B
aNSC enriched v.s. qNSC
translation
macromolecule biosynthetic process
cellular metabolic process
cell population proliferation
system development
0
10
20
30
40
negative log10(p value)

**Figure S6: Cluster proportion of all ten cell clusters**

(A) UMAP projections separating NTG and 3xTG-AD samples for all ten cell clusters.
(B) The proportion of each cluster was presented separately for NTG and 3xTG-AD, as percentage of total cell number from ten clusters.
(C) Proportional difference between NTG and 3xTG-AD cluster proportions from B. Left are cell clusters with greater proportion in NTG and right are cell clusters with greater proportion in 3xTG-AD.

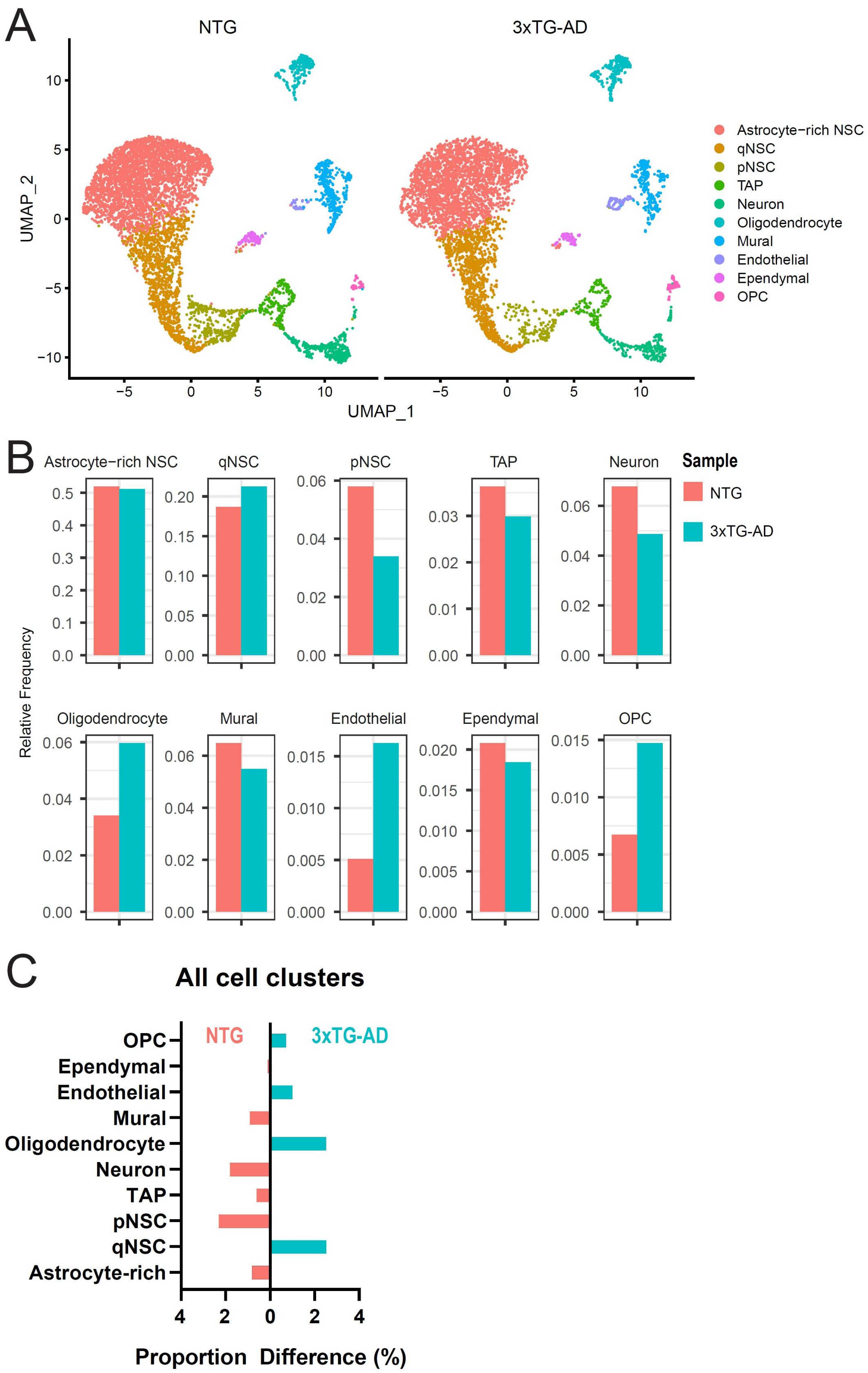
A
NTG
3xTG-AD
UMAP_2
UMAP_1
Astrocyte−rich NSC
qNSC
pNSC
TAP
Neuron
Oligodendrocyte
Mural
Endothelial
Ependymal
OPC
B
Astrocyte−rich NSC
qNSC
pNSC
TAP
Neuron
Sample
NTG
3xTG-AD
Relative Frequency
Oligodendrocyte
Mural
Endothelial
Ependymal
OPC
C
All cell clusters
NTG
3xTG-AD
OPC
Ependymal
Endothelial
Mural
Oligodendrocyte
Neuron
TAP
pNSC
qNSC
Astrocyte-rich
Proportion Difference (%)

**Figure S7: VlnpIot of selected genes**

Expression of representative genes were shown for each cluster as Vlnplot, separated by samples.

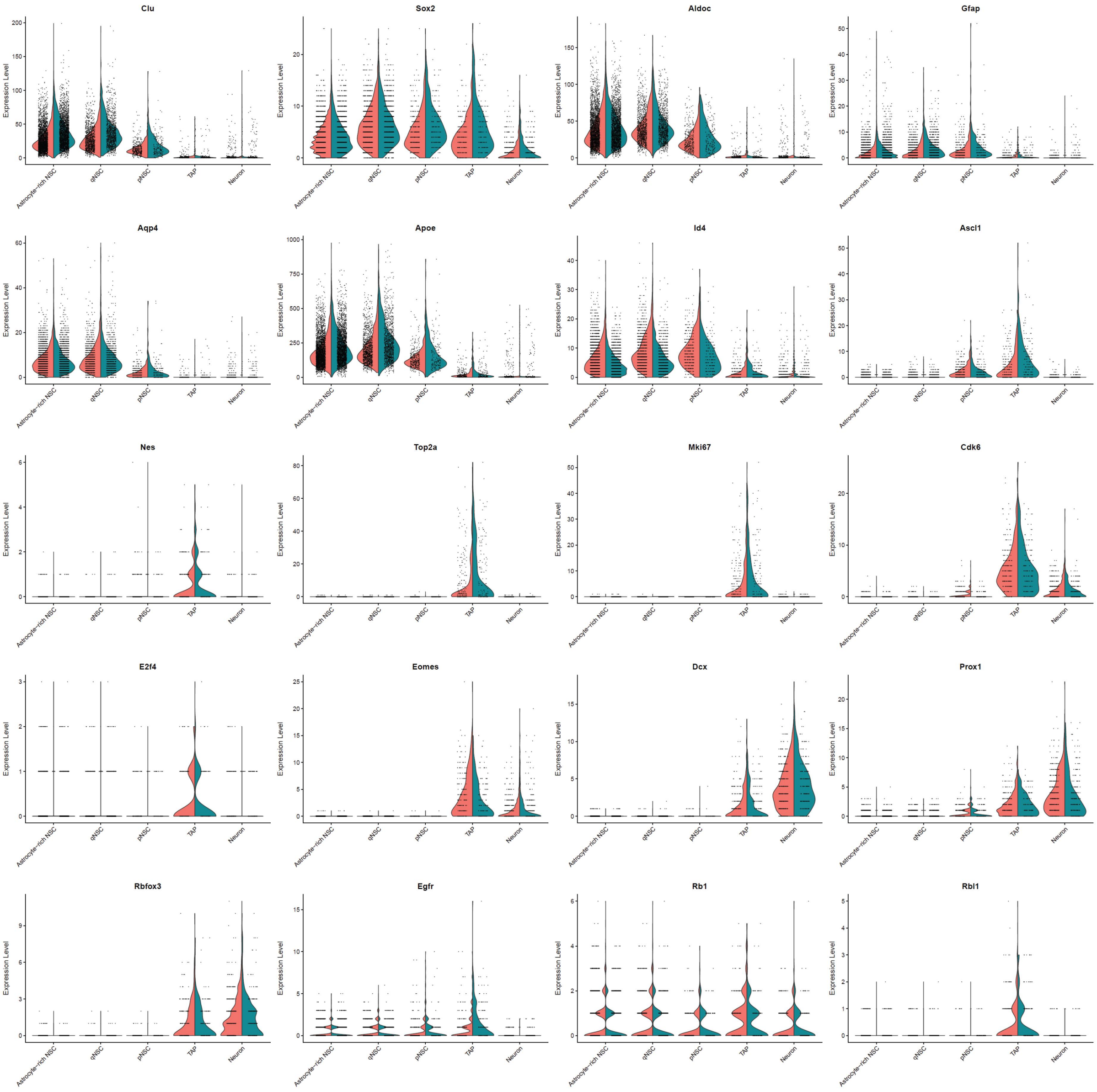
Clu
Sox2
Aldoc
Gfap
Aqp4
Apoe
Id4
Ascl1
Nes
Top2a
Mki67
Cdk6
E2f4
Eomes
Dcx
Prox1
Rbfox3
Egfr
Rb1
Rbl1
Expression Level
Astrocyte-rich NSC
qNSC
pNSC
TAP
Neuron

**Figure S8: Gene ontology analysis of differentially expressed genes in 3xTG-AD**

For each cell cluster, the up- or down-regulated genes were analyzed for gene ontology analysis using DAVID Bioinformatics online tool.

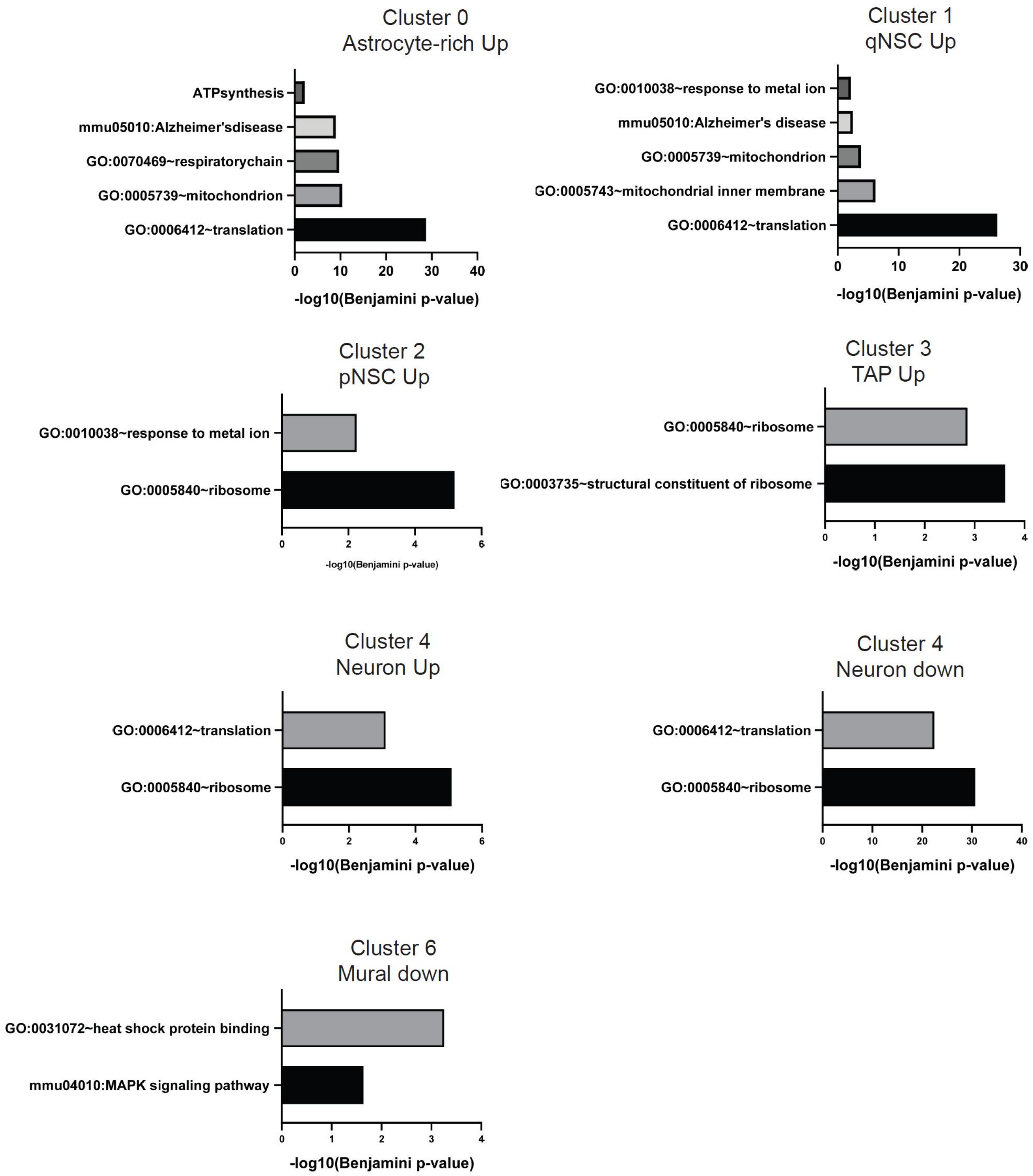
Cluster 0
Astrocyte-rich Up
ATPsynthesis
mmu05010:Alzheimer'sdisease
GO:0070469~respiratorychain
GO:0005739~mitochondrion
GO:0006412~translation
0 10 20 30 40
-log10(Benjamini p-value)
Cluster 1
qNSC Up
GO:0010038~response to metal ion
mmu05010:Alzheimer's disease
GO:0005739~mitochondrion
GO:0005743~mitochondrial inner membrane
GO:0006412~translation
0 10 20 30
-log10(Benjamini p-value)
Cluster 2
pNSC Up
GO:0010038~response to metal ion
GO:0005840~ribosome
0 2 4 6
-log10(Benjamini p-value)
Cluster 3
TAP Up
GO:0005840~ribosome
GO:0003735~structural constituent of ribosome
0 1 2 3 4
-log10(Benjamini p-value)
Cluster 4
Neuron Up
GO:0006412~translation
GO:0005840~ribosome
0 2 4 6
-log10(Benjamini p-value)
Cluster 4
Neuron down
GO:0006412~translation
GO:0005840~ribosome
0 10 20 30 40
-log10(Benjamini p-value)
Cluster 6
Mural down
GO:0031072~heat shock protein binding
mmu04010:MAPK signaling pathway
0 1 2 3 4
-log10(Benjamini p-value)

**Figure S9: RNA velocity and RNA expression plots for neural progenitor markers**

(A) RNA velocity of NTG and 3xTG-AD cells in "*cell*" mode.
(B) RNA velocity of NTG and 3xTG-AD cells in "*grid*" mode.
(C) Vlnplot for Hopx, Fabp7 (BLBP), Lpar1 and Ascl1.
(D) RNA velocity and RNA expression plots for Hopx, Fabp7 (BLBP), Lpar1 and Ascl1.

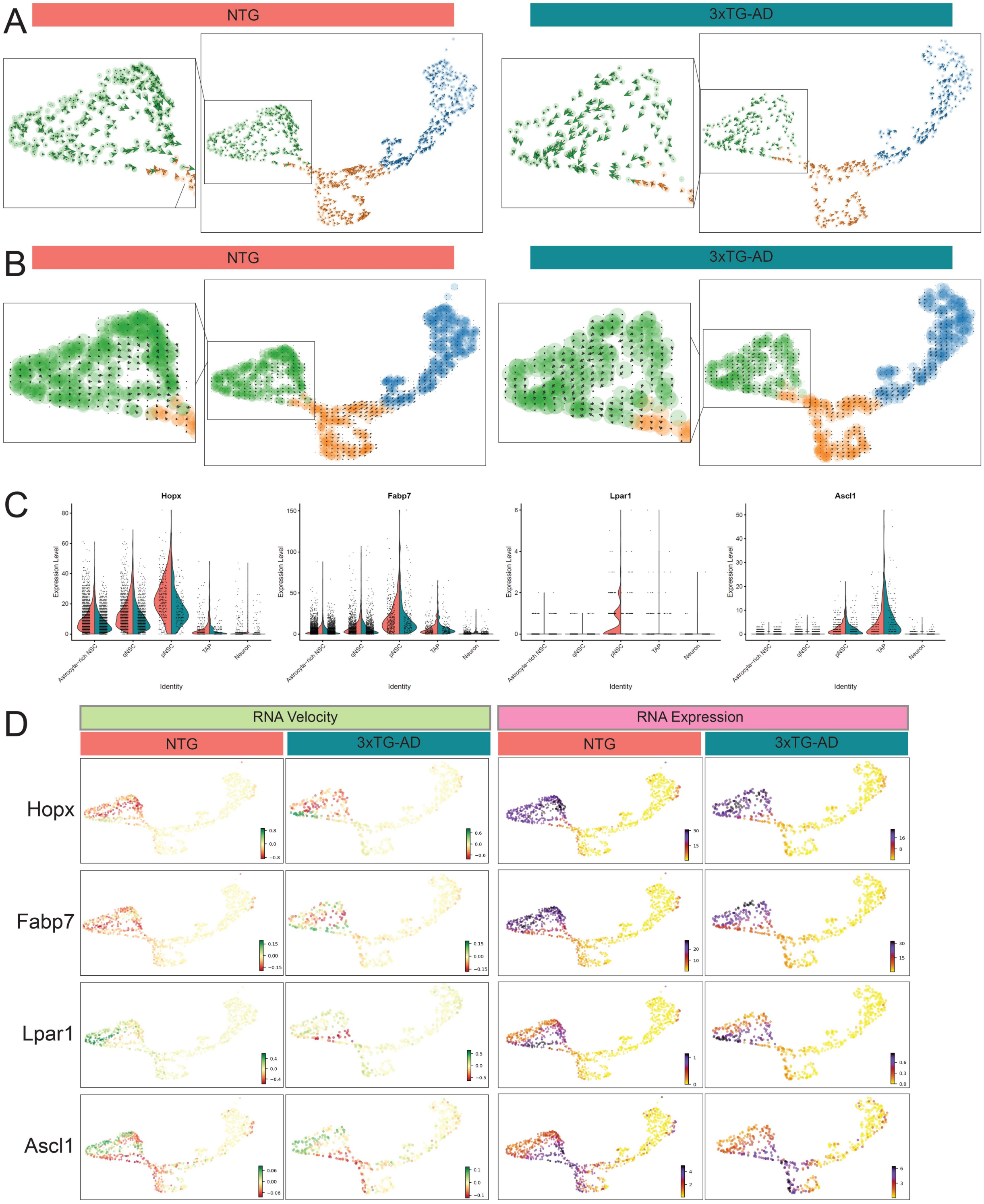
A
NTG
3xTG-AD
B
NTG
3xTG-AD
C
Hopx
Fabp7
Lpar1
Ascl1
Expression Level
Identity
Astrocyte-rich NSC
qNSC
pNSC
TAP
Neuron
D
RNA Velocity
RNA Expression
NTG
3xTG-AD
Hopx
Fabp7
Lpar1
Ascl1
